## Supplementary Information for "Transient power-law behaviour following induction distinguishes between competing models of stochastic gene expression"

**3. Short-time analysis for the variance and Fano factor of the mRNA counts** **8**

**4. Generalization of results to a broader class of systems with reversible reactions and gene state switching upon transcription** **9**

**5. Details of calculations performed to generate the main text figures** **13**

**References** **20**

**Supplementary Tables** **21**

**Supplementary Figures** **25**

S2 Finite sample size effects on estimating the short-time exponent using linear regression ($N = 5, j = 4$). 26
S3 Finite sample size effects on estimating the short-time exponent using non-linear regression ($N = 5, j = 4$). 27
S4 Finite sample size effects on estimating the short-time exponent using linear regression ($N = 5, j = 1$). 28
S5 Finite sample size effects on estimating the short-time exponent using non-linear regression ($N = 5, j = 1$). 29
S6 Effects of mRNA capture probability on estimating the short-time exponent using linear regression ($N = 5, j = 4$). . . . . . . 30
S7 Effects of mRNA capture probability on estimating the short-time exponent using non-linear regression ($N = 5, j = 4$). . . . . . . 31
S8 Effects of mRNA capture probability on estimating the short-time exponent using linear regression ($N = 5, j = 1$). . . . . . . 32
S9 Effects of mRNA capture probability on estimating the short-time exponent using non-linear regression ($N = 5, j = 1$). . . . . . . 33
S10 Choosing the optimal number of data points for the yeast data. . . . . . . 34
S11 Sampling distributions of parameters estimated from non-linear regression of the yeast data. . . . . . . 35
S12 Comparing fitting models with and without fixed time delays before transcription initiation. . . . . . . 36
S13 Estimation of the short-time exponent from mouse data. . . . . . . 37
S14 Comparing similar steady-state mRNA count distributions for models with 2, 3, and 4 inactive states to the effective telegraph model. . . . . . . 38
S15 Effects of varying the initial gene state distribution on the short-time behaviour of the mean mRNA count following induction. . . . . . . 39
S16 Estimation of the short-time exponent for different initial conditions and rate parameters. . . . . . . 40

### SUPPLEMENTARY NOTE 1. Mapping between the rate parameters of the telegraph model and the $N$-state model

In this Supplementary Note we derive the rate parameters of an effective telegraph model by writing these in terms of the rate parameters of the $N$-state model whose reaction scheme is given by Eq. (1) in the main text. We consider a parameter mapping involving the waiting time distribution between successive mRNA synthesis events.

##### 1.1 Moments of the waiting time distribution

Let $\tau$ denote the waiting time between successive mRNA synthesis events in the $N$-state model, and $f_N(\tau)$ its corresponding probability density function. The moments of $f_N(\tau)$ follow directly from the definition of the Laplace transform of $f_N(\tau)$, denoted by $f_N^*(s)$, since

$$f_N^*(s) = \langle e^{-s\tau}\rangle_N = \sum_{n=0}^{\infty}(-1)^n s^n \frac{\langle \tau^n\rangle_N}{n!}. \tag{1}$$

Suppose that the gene is in the active state $N$ at $t = 0$. Let $w_{i,j}(t)$ denote the mixed joint probability density function that the gene spends time $t$ in state $i$ before moving to state $j$. The probability distribution function of the total time to move from state $N$ to 1, and then back to state $N$ through states $2, 3, \ldots, N-1$ is the convolution of the individual distributions $w_{N,1}(t)$,$w_{1,2}(t)$,...,$w_{N-1,N}(t)$. Equivalently, the Laplace transform of the probability density function of the total time is given by

$$w^*_{N,1}(s)w^*_{1,2}(s)\cdot\dots\cdot w^*_{N-1,N}(s), \tag{2}$$

where $w^*_{i,j}(s)$ denote the Laplace transform of $w_{i,j}(t)$. Since the mRNA is transcribed only with some probability once the state $N$ is reached, it follows that any number $n \geq 0$ of the cycles ($N$ to 1 to 2 ... to $N$) is permitted before the mRNA is synthesised. This implies that the Laplace transform of the waiting time distribution between two successive mRNA synthesis events is given by

$$f_N^*(s) = \sum_{n=0}^{\infty}[w^*_{N,1}(s)w^*_{1,2}(s)\cdot\dots\cdot w^*_{N-1,N}(s)]^n w^*_{N,N}(s) = \frac{w^*_{N,N}(s)}{1 - w^*_{N,1}(s)w^*_{1,2}(s)\cdot\dots\cdot w^*_{N-1,N}(s)}, \tag{3}$$

where

$$w^*_{i,i+1}(s) = \frac{k_i}{s+k_i}, \quad i = 1,\ldots,N-1, \quad w^*_{N,1}(s) = \frac{k_N}{s+k_N+\rho}, \quad w^*_{N,N}(s) = \frac{\rho}{s+k_N+\rho}. \tag{4}$$

Note that $w^*_{N,N}(s)$ is the Laplace transform of the distribution of the time spent in state $N$ before an mRNA transcription event occurs. We also note that the series in Eq. (3) is guaranteed to converge since $w^*_{N,1}(s)w^*_{1,2}(s)\cdot\dots\cdot w^*_{N-1,N}(s) < 1$ for non-negative $s$. Substituting Eq. (4) into (3) gives

$$f_N^*(s) = \frac{\rho\prod_{i=1}^{N-1}(s+k_i)}{(s+k_N+\rho)\prod_{i=1}^{N-1}(s+k_i) - \prod_{i=1}^{N}k_i}. \tag{5}$$

This expression can be inverted to get $f_N(t)$ using partial fraction decomposition. The value of $f_N(0)$, however, can be obtained directly from $f_N^*(s)$ using the initial value theorem, yielding

$$f_N(0) = \lim_{s\to\infty}[sf_N^*(s)] = \rho. \tag{6}$$

From the expression for $f_N^*(s)$ and using Eq. (1), we get for the first two (raw) moments,

$$\langle\tau\rangle_N = -\left.\frac{df_N^*}{ds}\right|_{s=0} = \frac{k_N}{\rho}S_{N,1}, \tag{7}$$

$$\langle\tau^2\rangle_N = \left.\frac{d^2 f_N^*}{ds^2}\right|_{s=0} = 2\left[\left(1+\frac{k_N}{\rho}\right)S_{N,1} - \frac{1}{k_N}\right]\frac{k_N}{\rho}S_{N,1} - \frac{k_N}{\rho}(S_{N,1}^2 - S_{N,2}), \tag{8}$$

where

$$S_{N,1} = \sum_{j=1}^{N} \frac{1}{k_j}, \quad S_{N,2} = \sum_{j=1}^{N} \frac{1}{k_j^2}. \tag{9}$$

From here we find that the variance $\sigma_\tau^2 = \langle \tau^2 \rangle_N - \langle \tau \rangle_N^2$ is given by

$$\sigma_{\tau,N}^2 = \langle \tau^2 \rangle_N - \langle \tau \rangle_N^2 = \frac{k_N}{\rho} S_{N,2} + \frac{k_N}{\rho} \left(1 + \frac{k_N}{\rho}\right) S_{N,1}^2 - \frac{2}{\rho} S_{N,1}. \tag{10}$$

We note that the inverse of $\langle \tau \rangle_N$, which is the effective transcription rate, can be interpreted as the actual transcription rate $\rho$ multiplied by the steady-state probability that the gene is in the $N$th state,

$$P_{\mathrm{on},N} = \frac{1}{k_N \sum_{i=1}^{N} \frac{1}{k_i}}. \tag{11}$$

##### 1.2 The mRNA count distribution

The probability distribution of mRNA counts in steady-state conditions can be computed by laborious calculations using the chemical master equation (CME). Alternatively this can be more easily computed using results from queueing theory [1]. Namely, the $N$-state model maps to a queueing system with interarrival time $\tau$ whose probability density function is $f_N(\tau)$, and the service time is exponentially distributed with a rate parameter $d$. This queueing system belongs to the $G/M/\infty$ class of queueing systems, where $G$ stands for general or arbitrary interarrival time distribution, $M$ stands for Markovian or exponential service, and there are infinitely many servers. For this class of queueing systems the steady state distribution of the queue length (or in this case, the mRNA count) is known explicitly [2] (see also Ref. [1] for a recent review), and is given by

$$P(m) = \begin{cases} 1 - \sum_{j=1}^{\infty} (-1)^{j-1} \frac{C_{j-1}}{\langle \tau \rangle_N d j}, & m = 0 \\ \sum_{j=m}^{\infty} (-1)^{j-m} \binom{j}{m} \frac{C_{j-1}}{\langle \tau \rangle_N d j}, & m \geq 1 \end{cases}, \tag{12}$$

where $C_i$ for $i = 0, 1, \ldots$ are defined as

$$C_0 = 1, \quad C_j = \prod_{i=1}^{j} \frac{f_N^*(id)}{1 - f_N^*(id)}, \quad j \geq 1. \tag{13}$$

Note that since by Eq. (5) $f_N^*(s)$ is invariant to permutations of $k_i$, $i = 1, .., N-1$, it follows from Eq. (12) that the steady-state distribution of mRNA counts also shares this property.

We can further simplify Eq. (12) as follows. The probability-generating function $G(z)$ is given by

$$G(z) = \sum_{m=0}^{\infty} P(m) z^m = 1 + \sum_{m=1}^{\infty} \frac{C_{m-1}}{\langle \tau \rangle_N d m} (z-1)^m. \tag{14}$$

We now write the ratio $f_N^*(s)/[1 - f_N^*(s)]$ in the factorized form

$$\frac{f_N^*(s)}{1 - f_N^*(s)} = \frac{\rho \prod_{i=1}^{N-1} (s + k_i)}{\prod_{i=1}^{N} (s + k_i) - \prod_{i=1}^{N} k_i} = \frac{\rho (s + k_1) \ldots (s + k_{N-1})}{s (s + b_1), \ldots (s + b_{N-1})}, \tag{15}$$

where $b_1, \ldots, b_{N-1}$ are the non-zero roots of the polynomial in the denominator. If $N \leq 5$, then all $b_i$ can be found explicitly, otherwise they must be computed numerically. Substituting Eq. (15) into (14), and recognizing that

$$\langle \tau \rangle_N = \frac{1}{\rho} \left( \frac{b_1 \ldots b_{N-1}}{k_1 \ldots k_{N-1}} \right), \tag{16}$$

we get the following expression for the probability generating function $G(z)$,

$$G(z) = {}_{N-1}F_{N-1}\left(k'_1,\ldots,k'_{N-1};b'_1,\ldots,b'_{N-1};\rho'\,(z-1)\right), \tag{17}$$

where $k'_i = k_i/d$ and $b'_i = b_i/d$ for $i = 1,\ldots,N-1$, $\rho' = \rho/d$, and ${}_pF_q(a_1,\ldots,a_p;b_1,\ldots,b_q;z)$ is the generalized hypergeometric function defined as

$${}_pF_q(a_1,\ldots,a_p;b_1,\ldots,b_q;z) = \sum_{m=0}^{\infty}\frac{(a_1)_m\ldots(a_p)_m}{(b_1)_m\ldots(b_q)_m}\frac{z^m}{m!}, \tag{18}$$

and $(x)_n = x(x+1)\ldots(x+n-1)$ is the rising factorial. The probability $P(m)$ is obtained by taking the $m$th derivative of $G(z)$ evaluated at $z=0$, which yields

$$P(m) = \frac{(\rho')^m}{m!}\frac{(k'_1)_m\ldots(k'_{N-1})_m}{(b'_1)_m\ldots(b'_{N-1})_m}\,{}_{N-1}F_{N-1}\left(k'_1+m,\ldots,k'_{N-1}+m;b'_1+m,\ldots,b'_{N-1}+m;-\rho'\right). \tag{19}$$

Note that all rate parameters in Eq. (19) are normalised by the degradation rate parameter $d$ and hence the steady-state distribution is only a function of non-dimensional parameters.

The moments of the mRNA count distribution can be obtained using the probability generating function $G(z)$ in Eq. (17). For example, for the steady-state mean mRNA count we get

$$\langle m\rangle = \left.\frac{dG}{dz}\right|_{z=1} = \rho'\left(\frac{k'_1\ldots k'_{N-1}}{b'_1\ldots b'_{N-1}}\right) = \frac{1}{\langle\tau\rangle_N d} = \frac{\rho}{k_N S_{N,1} d} = \frac{\rho}{k_N d\sum_{i=1}^{N}\frac{1}{k_i}}. \tag{20}$$

##### 1.3 Rate parameters of an effective telegraph model via moment matching

We now consider the case of $N=2$ (the telegraph model). Let $x = k_2/k_1$. Then from Eqs. (6), (7) and (10) we have

$$f_2(0) = \rho, \quad \langle\tau\rangle_2 = \frac{1+x}{\rho}, \quad \sigma^2_{\tau,2} = \frac{2x^2}{k_2\rho} + \frac{(x+1)^2}{\rho^2}. \tag{21}$$

To map the $N$-state model to the effective 2-state telegraph model, we seek $\rho^*$, $k_1^*$ and $k_2^*$ of the telegraph model such that the following waiting time statistics of the two models match:

$$f_2(0) = f_N(0), \quad \langle\tau\rangle_2 = \langle\tau\rangle_N, \quad \sigma^2_{\tau,2} = \sigma^2_{\tau,N}. \tag{22}$$

From these equations we get the following expressions for $\rho^*$, $k_1^*$ and $k_2^*$,

$$\rho^* = \rho, \quad k_1^* = \frac{2\left(\langle\tau\rangle_N - \frac{1}{\rho}\right)}{\sigma^2_{\tau,N} - \langle\tau\rangle_N^2}, \quad k_2^* = \frac{2\rho\left(\langle\tau\rangle_N - \frac{1}{\rho}\right)^2}{\sigma^2_{\tau,N} - \langle\tau\rangle_N^2}. \tag{23}$$

In this mapping, the effective rate parameters depend only on $\rho$, $k_1,\ldots,k_N$ but not on the degradation rate $d$. It can also be shown that this mapping guarantees that the mean mRNA number is also the same between the models. We also note that both $k_1^*$ and $k_2^*$ are always positive since $\langle\tau\rangle_N > 1/\rho$ and $\sigma^2_{\tau,N} > \langle\tau\rangle_N^2$, which can be verified from expressions in Eqs. (7) and (10).

The mapping in Eq. (23) can be interpreted by introducing the random time $\tau_{1\to N}$ it takes the gene to go from state 1 to state $N$. Since this time is a sum of independent exponentially distributed random variables with rate parameters $k_1,\ldots,k_{N-1}$, its mean and variance are given by

$$\langle\tau_{1\to N}\rangle = \sum_{j=1}^{N-1}\frac{1}{k_j}, \quad \langle\tau_{1\to N}^2\rangle - \langle\tau_{1\to N}\rangle^2 = \sum_{j=1}^{N-1}\frac{1}{k_j^2}. \tag{24}$$

Substituting these expressions into Eq. (23), we get

$$k_1^* = \frac{1}{\langle\tau_{1\to N}\rangle}\cdot\frac{2}{\mathrm{CV}^2_{1\to N}+1}, \quad k_2^* = k_N\cdot\frac{2}{\mathrm{CV}^2_{1\to N}+1}, \tag{25}$$

where $\mathrm{CV}_{1\to N}$ is the coefficient of variation of $\tau_{1\to N}$,

$$\mathrm{CV}^2_{1\to N} = \frac{\langle\tau_{1\to N}^2\rangle - \langle\tau_{1\to N}\rangle^2}{\langle\tau_{1\to N}\rangle^2} = \sum_{j=1}^{N-1}\frac{1}{k_j^2}\left(\sum_{j=1}^{N-1}\frac{1}{k_j}\right)^{-2}. \tag{26}$$

#### SUPPLEMENTARY NOTE 2. Short-time analysis for the mean mRNA count

Assuming Markovian dynamics, the CME for the $N$-state reaction system described by Eq. (1) in the main text reads:

$$\frac{d}{dt}P_1(n,t) = d(n+1)P_1(n+1,t) - dnP_1(n,t) + k_N P_N(n,t) - k_1 P_1(n,t),$$
$$\frac{d}{dt}P_i(n,t) = d(n+1)P_i(n+1,t) - dnP_i(n,t) + k_{i-1}P_{i-1}(n,t) - k_i P_i(n,t), \quad i \in \{2, N-1\}$$
$$\frac{d}{dt}P_N(n,t) = \rho(P_N(n-1,t) - P_N(n,t)) + d(n+1)P_N(n+1,t) - dnP_N(n,t) + k_{N-1}P_{N-1}(n,t) - k_N P_N(n,t), \quad (27)$$

where $P_i(n,t)$ is the probability of observing $n$ mRNA molecules when the gene is in state $i$ at time $t$.

We first derive the moment equations from the CME. For compactness and simplicity of the expressions, we non-dimensionalise time and the reaction rates by the mRNA degradation rate: $t \mapsto td$, $k_i \mapsto k_i/d$ and $\rho \mapsto \rho/d$. This convention is used in this Supplementary Note and all the ones that follow (unless otherwise stated).

The $r^{th}$ moment of the mRNA, conditional on the gene being in state $i$, is given by:

$$(n^r)_i = \sum_n n^r P_i(n). \quad (28)$$

Note that the case $r = 0$ gives the probability of being in gene state $i$. In what follows, unless otherwise stated, for notational convenience we suppress the time dependence of the probabilities and of the moments.

By summing the terms on the right and left hand sides of Eq. (27) over $n$, we find the time-evolution equations for the probabilities of being in each gene state:

$$\frac{d}{dt}(n^0)_1 = k_N(n^0)_N - k_1(n^0)_1, \quad (29)$$
$$\frac{d}{dt}(n^0)_i = k_{i-1}(n^0)_{i-1} - k_i(n^0)_i, \quad i \in \{2, N\}. \quad (30)$$

Similarly by multiplying the right and left hand sides of Eq. (27) by $n$ and summing over $n$, we obtain the time-evolution equations for the first moments of mRNA counts (conditional on the gene state):

$$\frac{d}{dt}(n)_1 = -(n)_1 + k_N(n)_N - k_1(n)_1,$$
$$\frac{d}{dt}(n)_i = -(n)_i + k_{i-1}(n)_{i-1} - k_i(n)_i, \quad i \in \{2, N-1\},$$
$$\frac{d}{dt}(n)_N = \rho(n^0)_N - (n)_N + k_{N-1}(n)_{N-1} - k_N(n)_N. \quad (31)$$

Say the gene is initially in inactive state $j$ where $j \in 1,..,N-1$ and that the number of mRNA is zero. Due to the uni-directional nature of the gene state transitioning processes, the only reaction that can occur for very short time times is $G_j \to G_{j+1}$. Since at $t = 0$, $(n^0)_j = 1$ and $(n^0)_i = 0$, $\forall i \neq j$, it immediately follows from Eq. (30) that for short times $(n^0)_{j+1} \approx k_j t$.

The time-evolution for the next state to be visited, i.e. $j+2$, is given by

$$\frac{d}{dt}(n^0)_{j+2} = k_{j+1}(n^0)_{j+1} - k_{j+2}(n^0)_{j+2}. \quad (32)$$

Assuming a Taylor series expansion of $(n^0)_{j+2}$ in powers of $t$, integrating the above differential equation and using the previous result for $(n^0)_{j+1}$ it follows that to leading order, $(n^0)_{j+2} \approx \frac{1}{2}k_j k_{j+1} t^2$.

By iterating this procedure, one finds that generally

$$(n^0)_{j+r} \approx \frac{\prod_{i=j}^{j+r-1} k_i}{r!} t^r, \quad r = 1, ..., N-j. \quad (33)$$

Now summing Eq. (31), we obtain a time-evolution equation for the total mRNA count:

$$\frac{d}{dt}\langle m \rangle = \rho(n^0)_N - \langle m \rangle, \quad (34)$$

where $\langle m \rangle = \sum_{i=1}^{n} (n)_i$. Assuming a Taylor series expansion of $\langle m \rangle$ in powers of $t$, integrating the above differential equation and using Eq. (33), we finally obtain:

$$\langle m \rangle \approx \rho \frac{\prod_{i=j}^{N-1} k_i}{(N-j+1)!} t^{N-j+1}. \tag{35}$$

Next we derive an estimate of how small the time should be such that Eq. (35) holds. We will do this by deriving the next-to-leading order correction to the mean and then comparing the size of this term with the leading-order term previously derived.

From Eq. (30) it follows that

$$\frac{d}{dt}(n^0)_{j+r} = k_{j-1+r}(n^0)_{j-1+r} - k_{j+r}(n^0)_{j+r}, \quad j+r \in \{2, N\}. \tag{36}$$

Substituting $(n^0)_{j+r} = \alpha_{j,r} t^r + \beta_{j,r} t^{r+1} + O(t^{r+2})$ and collecting terms of order $t^r$, we obtain:

$$\beta_{j,r+1} = \frac{1}{r+2}(k_{j+r}\beta_{j,r} - k_{j+r+1}\alpha_{j,r+1}). \tag{37}$$

Note that $\alpha_{j,r}$ is known; by Eq. (33), it immediately follows that it is given by

$$\alpha_{j,r} = \frac{\prod_{i=j}^{j+r-1} k_i}{r!}. \tag{38}$$

Eq. (37) is a first-order non-homogeneous recurrence relation with variable coefficients. This can be solved using standard methods yielding:

$$\beta_{j,r} = \left[\prod_{i=0}^{r-1} f_i\right] \left(\beta_{j,0} + \sum_{m=0}^{r-1} \frac{g_m}{\prod_{i=0}^{m} f_i}\right), \tag{39}$$

where

$$f_r = \frac{k_{j+r}}{r+2}, \; g_r = \frac{-k_{j+r+1}\alpha_{j,r+1}}{r+2}. \tag{40}$$

Furthermore from Eqs. (29) and (30) and the initial conditions $(n^0)_j = 1$, and $(n^0)_i = 0, \;\; \forall i \neq j$, it follows that

$$(n^0)_j = 1 - k_j t + O(t^2),$$

which implies that

$$\beta_{j,0} = -k_j. \tag{41}$$

Hence from Eqs. (38), (39), (40) and (41) it follows that

$$\begin{aligned} (n^0)_N &= \alpha_{j,N-j} t^{N-j} + \beta_{j,N-j} t^{N-j+1} + O(t^{N-j+2}), \\ &= \frac{\prod_{i=j}^{N-1} k_i}{(N-j)!} t^{N-j} - \frac{\prod_{i=j}^{N-1} k_i}{(N-j+1)!} \left(\sum_{i=j}^{N} k_i\right) t^{N-j+1} + O(t^{N-j+2}). \end{aligned} \tag{42}$$

Finally substituting this result in the time-evolution for the total mRNA, Eq. (34), and computing the series solution we obtain:

$$\langle m \rangle = \rho \left[\prod_{i=j}^{N-1} k_i\right] \left[\frac{t^{N-j+1}}{(N-j+1)!} - (1 + \sum_{i=j}^{N} k_i) \frac{t^{N-j+2}}{(N-j+2)!}\right] + O(t^{N-j+3}). \tag{43}$$

Comparing the magnitudes of the terms proportional to $t^{N-j+1}$ and $t^{N-j+2}$, we find that the leading-order term, i.e. Eq. (35), is accurate provided

$$t \ll \frac{N-j+2}{1 + \sum_{i=j}^{N} k_i} \leq N+1, \quad j \in \{1, N-1\}. \tag{44}$$

The result suggests that the leading-order power law is visible over a long period of time provided the number of gene states to be traversed from the the initial inactive state to the active state $(N-j)$ is not too small and provided the rates of hopping from one gene state to another (relative to the mRNA degradation rate) are not too large. Note that Eq. (44) does not involve $\rho$ which implies that increasing $\rho$ increases the max mRNA count reached at the end of the time interval over which the leading-order power law is valid, without affecting the length of this interval.

##### 2.1 Post-transcriptional processing of mRNA

We note that in the calculations above we have interpreted the mRNA ($M$) in the $N$-state model as being the total mRNA. In some experiments, it is possible to distinguish between spliced and unspliced mRNA [3] and between nuclear and cytoplasmic mRNA [4]. We can model post-processing by adding extra reactions to the scheme described by Eq. (1) in the main text. For example, if we interpret $M$ as being unspliced nuclear mRNA then we could add a first-order reaction $M \to M_s$ describing the synthesis of spliced nuclear mRNA $M_s$ with some rate $k_s$. Since for short times, $d/dt\langle m_s\rangle \propto k_s\langle m\rangle$, it immediately follows by integrating Eq. (35) that the mean spliced mRNA count will follow a power law with an exponent that is one larger than that for unspliced mRNA. By the same argument the mean cytoplasmic mRNA will follow a power law with exponent that is one more than that of nuclear mRNA. If we interpret the mRNA in the $N$-state model as the cytoplasmic mRNA, then one could add a processing step to describe its translation to protein; in that case the short-time power law exponent for the latter will be one more than that for the cytoplasmic mRNA. On the other hand, if the post-processing of $M$ to a species $M_s$ can be modelled by a deterministic delay reaction (with delay $T$) then $\langle m_s\rangle(t) = \langle m\rangle(t-T)$ and hence in this case the power law exponent is unaffected by processing.

##### 2.2 Mixed gene state initial conditions

Thus far we have assumed a Dirac delta distribution on the initial gene state $j$. If we relax this assumption and allow for a mixed initial distribution of gene states, then it is straightforward from Eq. (35) to deduce that the mean mRNA count for short times is approximately given by

$$\langle m\rangle \approx \sum_{j=1}^{N-1} P_{G_j}(0) \times \rho \frac{\prod_{i=j}^{N-1} k_i}{(N-j+1)!} t^{N-j+1}. \tag{45}$$

Note that $P_{G_j}(0)$ is the probability that the gene is initially in state $j$, a quantity that we previously called $(n^0)_j(0)$ (see Eq. (28)).

If $P_{G_{N-1}}(0) \neq 0$ then the leading order term in Eq. (45) is quadratic in time ($t^2$). Similarly if $P_{G_{N-1}}(0) = 0$ and $P_{G_{N-2}}(0) \neq 0$ then the leading order term would be cubic in time ($t^3$). Generally if $P_{G_i}(0) = 0$ for $i = j+1...N-1$ and $P_{G_j}(0) \neq 0$ then the leading order term in Eq. (45) is proportional to $t^{N-j+1}$. Regardless, the short-time exponent remains a lower bound on the total number of gene states ($N-j+1 \leq N$).

#### SUPPLEMENTARY NOTE 3. Short-time analysis for the variance and Fano factor of the mRNA counts

We next compute the perturbation analysis for the variance. The initial conditions are the same as before: gene is in inactive state $j$ and the number of mRNA (and hence its variance) is initially zero.

From the CME, by a similar procedure to how we found the first moments, one can show that the time-evolution equations for the second moments of mRNA counts (conditional on the gene state) are given by

$$\begin{aligned} \frac{d}{dt}(n^2)_1 &= (n)_1 - 2(n^2)_1 + k_N(n^2)_N - k_1(n^2)_1, \\ \frac{d}{dt}(n^2)_i &= (n)_i - 2(n^2)_i + k_{i-1}(n^2)_{i-1} - k_i(n^2)_i, \quad i \in \{2, N-1\}, \\ \frac{d}{dt}(n^2)_N &= \rho[2(n)_N + (n^0)_N] + (n)_N - 2(n^2)_N + k_{N-1}(n^2)_{N-1} - k_N(n^2)_N. \end{aligned} \tag{46}$$

The second moment of the total mRNA count is defined to be $\langle m^2\rangle = \sum_{i=1}^n (n^2)_i$. Using Eqs. (31) and (46), it is straightforward to derive a time-evolution equation for the second moment:

$$\frac{d}{dt}\langle m^2\rangle = \rho[2(n)_N + (n^0)_N] + \langle m\rangle - 2\langle m^2\rangle. \tag{47}$$

To obtain the short-time solution for $(n)_N$ we consider its time-evolution equation as given by the last equation of Eq. (31). Since $(n)_i = 0\ \forall i$ and the short-time solution for $(n^0)_N$ is given by Eq. (42), it follows that

$$(n)_N = \frac{\rho\alpha_{j,N-j}}{N-j+1} t^{N-j+1} + \frac{\rho}{N-j+2}\left(\beta_{j,N-j} - \frac{(1+k_N)\alpha_{j,N-j}}{N-j+1}\right) t^{N-j+2} + O(t^{N-j+3}). \tag{48}$$

Substituting the short-time solutions for $(n^0)_N$, $\langle m \rangle$ (given by Eqs. (42) and (43), respectively) and the short-time solution for $(n)_N$ (Eq. (48)) in the time-evolution equation for the second moment of mRNA counts (Eq. (47)), and integrating we find

$$\langle m^2 \rangle = \rho \left[ \prod_{i=j}^{N-1} k_i \right] \left[ \frac{t^{N-j+1}}{(N-j+1)!} + (2\rho - 1 - \sum_{i=j}^{N} k_i) \frac{t^{N-j+2}}{(N-j+2)!} \right] + O(t^{N-j+3}). \tag{49}$$

Now the variance of total counts is defined as $\sigma_m^2 = \langle m^2 \rangle - \langle m \rangle^2$. Considering Eqs. (43) and (49), we immediately see that to next-to-leading order terms, the variance has the same short-time solution as $\langle m^2 \rangle$, i.e. Eq. (49).

We note that the leading-order term in Eq. (49) is the same as in Eq. (43) – this implies that to leading-order, the expressions for the mean and the variance are precisely the same. However the next-to-leading order terms differ which implies:

$$\sigma_m^2 - \langle m \rangle = \frac{2\rho^2 \prod_{i=j}^{N-1} k_i}{(N-j+2)!} t^{N-j+2} + O(t^{N-j+3}). \tag{50}$$

Since the difference is positive, it follows that the variance deviates faster from the leading-order term than the mean. From Eq. (50), it also follows that the Fano factor (defined as the variance divided by the mean of the total mRNA counts) is given by:

$$FF_m = \frac{\sigma_m^2}{\langle m \rangle} = 1 + \frac{2\rho}{N-j+2} t + O(t^2). \tag{51}$$

#### SUPPLEMENTARY NOTE 4. Generalization of results to a broader class of systems with reversible reactions and gene state switching upon transcription

We consider a general $N$-state model described by the following reaction scheme:

$$G_i \underset{k_{i,j}}{\overset{k_{j,i}}{\rightleftharpoons}} G_j,\ i,j = 1,\ldots,N, \quad G_N \xrightarrow{\rho} G_K + M, \quad M \xrightarrow{d} \emptyset. \tag{52}$$

This model includes the $N$-state model presented in the main text, for which $k_{i,i+1} = k_i$ for $i = 1,\ldots,N-1$, $k_{N,1} = k_N$, $K = N$, and all other rates $k_{i,j}$ are equal to 0. We note that in this Appendix we keep the absolute values of $k_{i,j}$ and $\rho$, i.e. we do not rescale them or time by $d$.

We assume that the gene initially starts in state $I$. We denote the mRNA number at time $t$ by $m(t)$, and assume that there is initially no mRNA present, i.e. $m(0) = 0$. Under these conditions, we derive a power law for the mean number of mRNA for short times, in the form of

$$\langle m(t) \rangle = \frac{A}{n!} t^n + O(t^{n+1}). \tag{53}$$

We show that $A$ and $n$ are determined from the series expansion of the Laplace transform of the probability density function of the waiting time until the first mRNA synthesis event,

$$f_1^*(s) = \frac{A}{s^n} + O\left(\frac{1}{s^{n+1}}\right), \quad s \to \infty. \tag{54}$$

Furthermore, the exponent $n$ is equal to the minimal number of states that are visited from the initial state $I$ at time $t = 0$ until the synthesis of the first mRNA molecule. To derive these results, we use a mapping of the reaction scheme in Eq. (52) to a $G/M/\infty$ queueing system, where $G$ denotes that the arrival (of mRNA) constitute a renewal process with a general (arbitrary) interarrival time distribution, $M$ denotes that the service (mRNA degradation) time is exponentially distributed (Markovian or memoryless), and there are infinitely many servers (meaning that mRNA can be in principle degraded as soon as it synthesised). For more details on how gene expression models map to infinite-server queues, we refer the reader to a recent review in Ref. [1].

We denote by $f(t)$ the probability density function of the interarrival time between two successive mRNA synthesis events. The Laplace transform of $f(t)$ was derived in SI Note 1. for the $N$-state model described in the main text, see Eq. (5). For the general model described by Eq. (52), $f(t)$ can be computed by solving the master equation for the probability $P_i(t)$ to find the gene in state $i$ at time $t$,

$$\frac{dP_i}{dt} = \sum_{j\neq i} k_{j,i} P_j - \sum_{j\neq i} k_{i,j} P_i, \quad i = 1, \ldots, N-1, \tag{55}$$

$$\frac{dP_N}{dt} = \sum_{j\neq N} k_{j,N} P_j - \left( \sum_{j\neq i} k_{N,j} + \rho \right) P_N, \tag{56}$$

where the reaction $G_N \to G_K + M$ is treated as an absorbing state, and the initial condition is

$$P_i(0) = \delta_{i,K}, \quad i = 1, \ldots, N. \tag{57}$$

If we denote $\mathbf{P}(t) = (P_1(t), \ldots, P_N(t))$, then the master equation (55) can be written as

$$\frac{d\mathbf{P}}{dt} = \mathbf{P}D, \quad \mathbf{P}(0) = \mathbf{e}_K \equiv (\underbrace{0, \ldots, 0}_{K-1}, 1, \underbrace{0, \ldots, 0}_{N-K}), \tag{58}$$

where $D$ is $N \times N$ matrix whose matrix elements are defined as

$$D_{ij} = \begin{cases} k_{ij}, & i \neq j, \\ -\sum_{m\neq i} k_{i,m}, & i = j \neq N, \\ -\sum_{m\neq i} k_{i,m} - \rho, & i = j = N. \end{cases} \tag{59}$$

The solution of Eq. (58) is

$$\mathbf{P}(t) = \mathbf{e}_K e^{Dt}. \tag{60}$$

The probability density function of the waiting time between two successive mRNA synthesis events is given by

$$f(t) = \rho P_N(t) = \mathbf{e}_K e^{Dt}(-D\mathbf{1}^T), \tag{61}$$

where $\mathbf{1}^T$ is $N \times 1$ column vector with every element being equal to 1. This distribution is known as a phase-type distribution [5]. If the initial state $I \neq K$ (the state to which the gene switches after producing an mRNA molecule), the probability density function of the waiting time until the first mRNA synthesis event is not given by $f(t)$, but is instead given by $f_1(t)$, which reads

$$f_1(t) = \mathbf{e}_I e^{Dt}(-D\mathbf{1}^T), \quad \mathbf{e}_I = (\underbrace{0, \ldots, 0}_{I-1}, 1, \underbrace{0, \ldots, 0}_{N-I}). \tag{62}$$

The waiting time densities $f(t)$ and $f_1(t)$, the initial condition $m(0) = 0$ and the service rate $d$ fully specify the queueing system $G/M/\infty$, for which we are interested in finding the early-time behaviour of the mean mRNA number $\langle m(t) \rangle$. The case in which $f_1(t) = f(t)$, which in our model corresponds to the initial state $I = K$, has been solved in Ref. [2], yielding the following result for the Laplace transform of the mean mRNA number,

$$\mathcal{L}[\langle m(t) \rangle](s) = \frac{f^*(s)}{(s+d)[1 - f^*(s)]}. \tag{63}$$

Below we generalize this result to $f_1(t) \neq f(t)$, which yields

$$\mathcal{L}[\langle m(t) \rangle](s) = \frac{f_1^*(s)}{(s+d)[1 - f^*(s)]}. \tag{64}$$

We note that $f_1^*(s)$ is a rational function of $s$, i.e. it can be written as a ratio of two coprime polynomials $p(s)$ and $q(s)$,

$$f_1^*(s) = \frac{p(s)}{q(s)} = \frac{As^{\deg(p)} + \ldots}{s^{\deg.(q)} + \ldots}, \tag{65}$$

where we have written only the highest-order terms in each of the polynomials. To get an early-time behaviour of $\langle m(t)\rangle$, we expand $f_1^*(s)$ in $s$ as $s\to\infty$,

$$f_1^*(s) = \frac{A}{s^n} + O\left(\frac{1}{s^{n+1}}\right), \quad s\to\infty, \quad n = \deg(p) - \deg(q). \tag{66}$$

and substitute back into Eq. (64), obtaining

$$\mathcal{L}[\langle m\rangle](s) = \frac{A}{s^{n+1}} + O\left(\frac{1}{s^{n+2}}\right), \tag{67}$$

where we have used the fact that $f^*(s)\to 0$ for $s\to\infty$ (hence, there is no contribution from $f^*(s)$ to the lowest order). The power law in Eq. (53) follows from the inverse Laplace transform of $1/s^{n+1}$, which gives $t^n/n!$. According to Theorem 3.1 in Ref. [6], given a phase-type probability density function whose Laplace transform is $p(s)/q(s)$, where $p(s)$ and $q(s)$ are coprime polynomials, the difference $n = \deg(q) - \deg(p)$ is equal to the minimal number of states that are visited before the absorption (in our case the synthesis of mRNA). This completes the derivation of the power law in Eq. (53), and the interpretation of $A$ and $n$.

To derive Eq. (64), we reiterate the steps in Ref. [2], adjusted to the first waiting time described by $f_1(t)$ rather than $f(t)$. Let $T_n$ denotes the arrival time of the $n$th mRNA, and $\chi_n$ its degradation time. The number of mRNA molecules at time $t$ can be written as

$$m(t) = \sum_{T_n\le t} \theta(t-T_n, \chi_n), \tag{68}$$

where the function $\theta(u,x)$ is defined as

$$\theta(u,x) = \begin{cases} 1, & 0\le u\le x, \\ 0, & \text{otherwise.} \end{cases} \tag{69}$$

Conditioned on $T_1 = x$, $m(t)$ can be written as

$$m(t) = \begin{cases} \theta(t-T_1,\chi_1) + m_0(t-x), & x\le t, \\ 0, & x > t, \end{cases} \tag{70}$$

where $m_0(t)$ is the number of mRNA molecules at time $t$ in a process in which the time until the first mRNA synthesis event is distributed according to $f$ instead of $f_1$ (another way to say this is that time $t=0$ in this process is an arrival epoch). For $x\le t$, the conditional probability to find $m(t)=k$ mRNA molecules at time, given that the first arrival occurred at time $t=x$, is given by

$$P(m(t)=0|T_1=x) = P(m_0(t)=0)P(\chi_1\le t-x), \tag{71a}$$

$$P(m(t)=k|T_1=x) = P(m_0(t-x)=k)P(\chi_1\le t-x) + P(m_0(t-x)=k-1)P(\chi_1 > t-x), \quad k\ge 1, \tag{71b}$$

where

$$P(\chi_1\le t-x) = 1-e^{-d(t-x)}, \quad P(\chi_1 > t-x) = e^{-d(t-x)} \tag{72}$$

On the other hand, for $x > t$,

$$P(m(t)=k|T_1=x) = \delta_{k,0}. \tag{73}$$

We now define the probability generating function

$$G(z,t) = \sum_{k=0}^{\infty} P(m(t)=k)z^k = \sum_{k=0}^{\infty}\int_0^{\infty} dx\ f_1(x)P(m(t)=k|T_1=x)z^k. \tag{74}$$

Inserting Eqs. (71), (72) and (73) into Eq. (74) yields

$$G(z,t) = 1 - F_1(t) + \int_0^t dx\ f_1(x)G_0(z,t-x)\left[1+(z-1)e^{-d(t-x)}\right], \tag{75}$$

where

$$G_0(z,t) = \sum_{k=0}^{\infty} P(m_0(t) = k) z^k. \tag{76}$$

By taking the Laplace transform on both sides of Eq. (75), we get

$$G^*(z,s) = \frac{1 - f_1^*(s)}{s} + f_1^*(s) G_0^*(z,s) + (z-1) f_1^*(s) G_0^*(z, s+d), \tag{77}$$

where

$$G^*(z,s) = \int_0^{\infty} dt\, G(z,t) e^{-st}, \quad G_0^*(z,s) = \int_0^{\infty} dt\, G_0(z,t) e^{-st}. \tag{78}$$

Similarly, for the process in which the initial time $t = 0$ is an arrival epoch we get

$$G_0^*(z,s) = \frac{1 - f^*(s)}{s} + f^*(s) G_0^*(z,s) + (z-1) f^*(s) G_0^*(z, s+d). \tag{79}$$

By combining these two equations we get

$$G^*(s) = \frac{1}{s} + \frac{f_1^*(s)}{f^*(s)} \left[ G_0^*(z,s) - \frac{1}{s} \right]. \tag{80}$$

The Laplace transform of the mean mRNA number $\langle m(t) \rangle$ is given by

$$\mathcal{L}[\langle m(t) \rangle](s) = \left. \frac{dG}{dz} \right|_{z=1} = \frac{f_1^*(s)}{f^*(s)} \mathcal{L}[\langle m_0(t) \rangle](s) = \frac{f_1^*(s)}{(s+d)[1 - f^*(s)]}, \tag{81}$$

where in the last equality we used the expression for the Laplace transform of $\langle m_0(t) \rangle$ in Eq. (63).

Let us now apply this result to the $N$-state model in the main text, with state $j$ as the initial state as time $t = 0$. For $j \neq N$, The Laplace transform $f_1^*(s)$ reads

$$f_1^*(s) = \left( \prod_{i=j}^{N-1} \frac{k_i}{s + k_i} \right) f_N^*(s), \tag{82}$$

where $f_N^*(s)$ is given by Eq. (5). Expanding $f_1^*(s)$ in $s$ at $s \to \infty$, we get

$$f_1^*(s) = \frac{\rho \prod_{i=j}^{N-1} k_i}{s^{N-j+2}} + O\left( \frac{1}{s^{N-j+3}} \right), \tag{83}$$

from which we get that

$$A = \rho \prod_{i=j}^{N-1} k_i, \quad n = N - j + 1, \tag{84}$$

exactly as Eq. (3) in the main text.

So far we assumed that the initial mRNA count at $t = 0$ was zero. Let us now relax this assumption and assume that $m(0)$ follows some initial mRNA distribution which is not a delta function $\delta_{m(0),0}$. Let us denote by $m_{\text{init}}(t)$ the number of mRNA molecules present at time $t$ from the group of mRNA molecules that were present initially at time $t = 0$, so that $m_{\text{init}}(0) = m(0)$. We can then write $m(t)$ as a sum of two random variables, $m(t) - m_{\text{init}}(t)$ and $m_{\text{init}}(t)$. The first random variables is described by the previously discussed theory in which we assumed $m(0) = 0$. The second random variable follows a binomial distribution

$$P(m_{\text{init}}(t) = k) = \binom{m(0)}{k} e^{-dkt} \left( 1 - e^{-dt} \right)^{m(0)-k}, \tag{85}$$

where $e^{-dt}$ is the survival probability of a single mRNA molecule until time $t$. To derive this result, we note that: (1) mRNA degradation times are exponentially distributed, (2) the exponential distribution is memoryless, and (3) mRNA molecules are degraded independently of each other. From (1) and (2) it follows that the residual lifetimes of mRNA molecules that were present at time $t = 0$ follow the same exponential distribution as the degradation times. Therefore, the survival probability of a single mRNA molecule until time $t$ is equal to $e^{-dt}$. Together with (3), this means that the number of mRNA molecules that have survived until $t$ follows a binomial distribution with the success probability $e^{-dt}$. Putting everything together, we get

$$\langle m(t)\rangle - \langle m(0)\rangle e^{-dt} = \frac{A}{n!}t^n + O(t^{n+1}), \tag{86}$$

where $\langle m(0)\rangle e^{-dt}$ is the mean of the binomial distribution in Eq. (85) averaged over the initial mRNA distribution. Hence, the short-time power law is preserved, provided the exponential decay is substracted from the mean.

Finally, we can extend our initial model further, assuming that it takes some time for the inducer molecule to reach the gene. In that case, our model does not start in $G_1$ at $t = 0$, but instead starts in some unknown state $U_1$. The induction process then goes through a series of unknown states until it eventually reaches state $G_1$. If we assume that the number of these unknown states is large, then the time it takes the inducer to get from $U_1$ to $G_1$ is sharply peaked at some mean time $t_0$. This can be accounted for in our model by adding a deterministic delay to the time it takes a gene to produce the first mRNA molecule. In that case, the probability density $f_1(t)$ of the time until first mRNA molecules is produced is replaced by $f_1(t - t_0)$, whose Laplace transform is given by $f_1^*(s)e^{-st_0}$. This yields the following expression for the mean mRNA count $\langle m(t)\rangle$,

$$\langle m(t)\rangle - \langle m(0)\rangle e^{-dt} = \begin{cases} 0 & t < t_0, \\ \frac{A}{n!}(t - t_0)^n + O(t^{n+1}) & t \geq t_0. \end{cases} \tag{87}$$

Again, the power law remains the same, but the time is now shifted by the deterministic delay $t_0$.

#### SUPPLEMENTARY NOTE 5. Details of calculations performed to generate the main text figures

##### 5.1 Fig. 2 in the main text

The rate parameter values of the $N$-state model were sampled to sufficiently cover the parameter space suggested by experimental data. Next we define the effective gene activation rate ($\sigma_u$), gene inactivation rate ($\sigma_b$), and transcription rate ($\rho_u$) of the $N$-state model and choose them to lie approximately within the bounds estimated from experimental data (see Table 1 in [7]). This was done to reflect the diversity of observed gene expression signatures in a variety of eukaryotic cells.

For the $N$-state model, the effective gene activation rate is roughly given by the inverse of the time it takes to transition from gene state 1 to $N$, i.e. $\sigma_u = (\sum_{i=1}^{N-1} k_i^{-1})^{-1}$, the effective gene inactivation rate is simply the inverse of the time it takes to leave the active state $N$, i.e. $\sigma_b = k_N$ and the effective transcription rate $\rho_u = \rho$. Note that these formulae are only used to choose meaningful rate parameter ranges for the $N$-state model.

From Table 1 in [7], after normalising each of the estimated rate parameters by the mRNA degradation rate parameter, the minimum and maximum values were: 0.22-11.3 ($\sigma_u$), 3.18-243.6 ($\sigma_b$), and 28.9-6300 ($\rho_u$), respectively. Hence the $N$-state rate parameters were chosen such that the observed rate parameters $\sigma_u$, $\sigma_b$, and $\rho_u$ were log-uniformly distributed in the following ranges: $\sigma_u \in [0.025, 20]$, $\sigma_b \in [0.1, 250]$, and $\rho_u \in [10, 6500]$. Specifically $k_i$ was chosen to be log-uniformly distributed on $[0.1, 40]$; $k_N$ was chosen to be log-uniformly distributed on $[0.1, 250]$; and $\rho$ was chosen to be uniformly distributed on $[10, 6500]$, for each $N = 3, 4, 5$.

Steady-state distributions of mRNA counts were computed using the finite-state projection algorithm (FSP) [8]. Rate parameter sets which gave steady-state mean mRNA count $< 2$ and $> 200$ were filtered out, as these distributions are either: in the former case, trivial to approximate due to very low copy numbers; or, in the latter case, have excessively large copy numbers, beyond those of common experimental distributions. The maximum mRNA count to truncate the FSP state space was selected to be the minimum of either the steady-state mean plus 10 steady-state standard deviations or 800. The absolute minimum size of the state space was chosen to be 20 mRNA.

In order for summary statistics to be comparable between the $N = 3, 4, 5$ state models, rate parameter sets were sampled such that the effective sampling density was constant for all values of $N$. This effectively enforces an approximate equal spread of samples across the different $N$-state parameter spaces. If $L$ represents the total number of samples, $y$ the average number of samples per dimension (sampling density), and $q$ the dimensionality of the parameter space (this equals $N+1$ since we have $N$ gene state transition parameters $k_1, .., k_N$ and $\rho$; the degradation rate parameter $d$ is fixed to 1), then they are connected by the following equality: $y = L^{1/q}$. Therefore the total number of samples for each model ($L$) was approximately: 1,200 (3-state), 8,200 (4-state), and 38,000 (5-state); preserving a sampling density of approximately 6.0. These procedures were used to construct Fig. 2A-G and I-K.

The rate parameter values used in Fig. 2D-G for the 5-state model steady-state mRNA count distribution shapes are given in Table S1.

Fig. 2H used the same procedure as detailed previously, with one exception. Since shapes I and IV were sampled highly infrequently for each value of $N$, additional rate parameter sets were generated to ensure a sufficient sample size to draw summary statistics from. Note that these additional samples were not included in the results seen in Fig. 2I-K. For sampling parameters that resulted in distribution shape IV, $k_i$ was chosen to be uniformly distributed in [1, 4], for each $N = 3, 4, 5$, and $k_N$ was uniformly distributed on [0.1,1]. The sampling range of $\rho$ remained unchanged. Summary statistics for Fig. 2H are shown in Table S2.

##### 5.2 Fig. 3 in the main text

The reaction network used in Fig. 3A-C was:

$$\begin{aligned} &G_1 \xrightarrow{k_1} G_2 \xrightarrow{k_2} G_3 \xrightarrow{k_3} G_4 \xrightarrow{k_4} G_5 \xrightarrow{k_5} G_1, \\ &G_5 \xrightarrow{\rho} G_5 + M, \\ &M \xrightarrow{d} \emptyset. \end{aligned} \tag{88}$$

The reaction network used in Fig. 3D was:

$$\begin{aligned} &G_1 \xrightarrow{k_1} G_2 \xrightarrow{k_2} G_3 \xrightarrow{k_3} G_4 \xrightarrow{k_4} G_5 \xrightarrow{k_5} G_1, \\ &G_5 \xrightarrow{\rho} G_5 + M, \\ &M \xrightarrow{\gamma} M + P, \\ &M \xrightarrow{d} \emptyset, P \xrightarrow{d_p} \emptyset. \end{aligned} \tag{89}$$

The reaction networks used in Fig. 3F were the following. For the blue ($t^3$) line:

$$\begin{aligned} &G_1 \xrightarrow{k_1} G_2 \xrightarrow{k_2} G_3 \xrightarrow{k_3} G_4 \xrightarrow{k_4} G_5 \xrightarrow{k_5} G_1, \\ &G_2 \xrightarrow{k_{2,5}} G_5 \\ &G_5 \xrightarrow{\rho} G_5 + M, \\ &M \xrightarrow{d} \emptyset, \end{aligned} \tag{90}$$

and for the pink ($t^4$) line:

$$\begin{aligned} &G_1 \underset{k_{2,1}}{\overset{k_1}{\rightleftharpoons}} G_2 \xrightarrow{k_2} G_3 \xrightarrow{k_3} G_4 \xrightarrow{k_4} G_5 \xrightarrow{k_5} G_1, \\ &G_5 \xrightarrow{\rho} G_2 + M, \\ &M \xrightarrow{d} \emptyset. \end{aligned} \tag{91}$$

The rate parameter values used in Fig. 3A-D, and F are given in Table S3. The rate parameter values and gamma distribution parameters used in Fig. 3E for the static extrinsic noise SSA mean mRNA count results are given in Table S4. Calculations of the mean and variance were performed using DifferentialEquations.jl and MomentClosure.jl in Julia [9, 10].

##### 5.3 Fig. 4 in the main text

The rate parameters for the $N$-state gene model with $N = 2, 3, 4, 5$ were randomly chosen from the ranges $k_i = [0.1, 40.0]$ for $i = 1, .., N-1$, $k_N = [0.1, 250]$ and $\rho = [10, 6500]$. Note that these are the same ranges as used for Fig. 2 except that the ranges were sampled uniformly rather than log-uniformly. The rate degradation parameter $d$ is fixed to 1.

For each of these parameters, we use Eq. (3) in the main text to calculate the time at which the mean mRNA count should be 40 assuming that power law behavior dominates the time regime over which post-induction measurements are made. We call this time $t_{\text{on}}$. We chose 40 since this is the same order of magnitude as the median mRNA count per cell measured in exponentially growing mammalian cells [11]. If $t_{\text{on}}$ falls in the range $0.01 - 0.2$, we accept this parameter set and set $t_{\text{on}}$ as the largest time at which the mean mRNA is measured in our simulated induction experiment. Note that since the median lifetime of mRNA is 9 hours in mammalian cells [11] and time is non-dimensional (it is multiplied by the mRNA degradation rate), it follows that the range $t_{\text{on}} = 0.01 - 0.2$ corresponds to a real time range of approximately 5 minutes to 2 hours; this is a reasonably realistic time range to make several mRNA count measurements.

For each accepted parameter set, we then simulated the variation of the mean mRNA count over the time range $[0, t_{\text{on}}]$ by numerically integrating the time-evolution equations (given by the CME) for the mean mRNA count. Those parameter sets whose steady-state mean mRNA count was over 1000 or whose mean mRNA count at $t_{\text{on}}$ was less than 1 were removed, leaving us with a final total of 1,721,036 realistic parameter sets. For these, it was assumed that 5 equispaced measurements were made of the mRNA mean, i.e. at $t = t_{\text{on}}/5,\ 2t_{\text{on}}/5,\ 3t_{\text{on}}/5,\ 4t_{\text{on}}/5,\ t_{\text{on}}$. Linear regression on the log-log plot of mean counts versus time was used to estimate the exponent of the power law; this was performed using the "lm" function from the GLM.jl package in Julia. The distributions of the exponents calculated for all parameter sets of 10 configurations of the $N$-state model are shown in Fig. 4 B-H.

##### 5.4 Fig. 5 in the main text

A model of the form of Eq. (87) was fitted to the yeast data for Fig. 5, where $d$ is set to 0 (see main text for explanation). This implies that we need to estimate four parameters: $\langle m(0)\rangle, A, t_0$ and $n$.

Since there are simultaneous measurements of the nuclear and cytoplasmic mRNA for each gene (CTT1 and STL1) under each experimental condition (0.2 and 0.4 mol NaCl), the delay between the application of the stimulus and transcription initiation ($t_0$) would ideally be inferred using both mRNA curves. However due to the very small mean nuclear mRNA counts (mostly $< 1$ when power-law behaviour is evident) and resulting uncertainty in these small measurements, this delay is difficult to reliably infer using the nuclear mRNA measurements. Hence we instead fit the model to cytoplasmic mRNA data (which is much more abundant, mean $> 10$, when the power-law behaviour is evident) to infer the cytoplasmic mRNA parameters, including the delay, then fix this delay for the corresponding nuclear mRNA measurement and infer the remaining nuclear mRNA parameters.

The optimal number of time points to use in the regression was determined by fitting the model for each number of time points and choosing the maximum number of time points that minimised the residual sum of squares (RSS) between the data and model. This procedure is detailed in SI Note 6.4 and Fig. S10.

Data was taken from Ref. [4] and pre-processed by summing the nuclear (respectively cytoplasmic) mRNA count over all cells and dividing by the total number of cells to obtain the mean mRNA count across all repetitions of each experimental condition. Where NaN values were present in the original data, the mean mRNA at that time was computed with the remaining values. The number of cells varied for each of the eight time points and for the two different experimental conditions. The number of cells at each time point can be found in Table S6. In all cases at least 1000 cells were used to compute both the nuclear and cytoplasmic mean mRNA counts. Bootstrapping was used to generate a large number of mean versus time curves; fitting the model to each curve leads to a set of parameters, the sampling distributions of which are found in Fig. 5 in the main text and Fig. S11.

The model and corresponding RSS function were defined and passed to the "bboptimize" function from the BlackBoxOptim.jl package in Julia, along with the bounds on each parameter, to find the parameter set which minimises the RSS. This approach is equivalent to, and was used as an alternative to, non-linear least squares regression due to compatibility issues with the piecewise-defined model function and the non-linear regression tools in LsqFit.jl. The following bounds were used for each parameter: prefactor, $A \in [0.0, 10^3]$; exponent, $n \in [1.0, 9.0]$; delay, $t_0 \in [0.0, 5.0]$; and constant mean mRNA count during delay, $\langle m(0)\rangle \in [0.0, 1.0]$.

The error bars for each mean mRNA curve display the standard deviation of the sampling distribution of each mean mRNA count (i.e. the standard error of the mean, SEM), computed from 10,000 bootstrapped samples of the mean mRNA count at each time point. Due to the significant sample size at each time point, these error bars are quite small.

#### SUPPLEMENTARY NOTE 6. Details of calculations performed to generate the supplementary information figures

##### 6.1 Fig. S1 - Additional steady-state mRNA count distribution shapes

Fig. S1 shows various additional steady-state mRNA count distribution shapes for each of the shape classes I-IV described in Fig. 2D-G. Here we compare the distributions from the $N$-state model with $N = 5$ with those from the effective telegraph model. The corresponding rate parameters for the 12 plots can be found in Table S5.

Fig. S1C displays a notable distribution shape, representing a very slow-switching regime of the telegraph model where the timescales of gene activation and inactivation are much longer than those of mRNA degradation leading to a distribution that is well approximated by the weighted sum of a Dirac delta distribution at 0 mRNA (due to the inactive state) and a Poisson distribution (due to the active state). Fig. S1L displays the limiting case when the effective telegraph model gives a distribution with a long plateau starting at 0 molecules. This distribution is obtained when the parameters of the effective telegraph model are $k_1 = k_2 = d$ and $\rho \geq d$. Other notable shapes include Fig. S1F showing a left-skewed distribution with a long plateau, and Fig. S1J showing a distribution with high probability at very low copy numbers, representing mRNA that is degraded rapidly and/or a gene that is highly inactive.

##### 6.2 Figs. S2-S5 - Finite sample size effects on the estimation of the short-time exponent

To investigate finite sample size effects on the estimation of the short-time exponent we used the stochastic simulation algorithm (SSA) to simulate the mRNA count in a population of cells over time and computed the mean mRNA count over various different sample sizes. The sample sizes used were $100, 500, 1,000$ and $10,000$; chosen to cover the range of sample sizes reported in smFISH, single-cell RNAseq and bulk RNAseq experiments. We chose 20 different parameter sets from those generated according to the procedure detailed in SI Note 5. 3 for Fig. 4 in the main text for the modelling configurations: $N = 5, j = 4$; and $N = 5, j = 1$. These parameter sets represent mean mRNA counts varying between 1 and 25 by the time of the final sample. The mean mRNA count was simulated for a total time of $t_{\text{on}}$ and measured at the following times: $t = t_{\text{on}}/5,\ 2t_{\text{on}}/5,\ 3t_{\text{on}}/5,\ 4t_{\text{on}}/5,\ t_{\text{on}}$.

Both linear and non-linear regression techniques were applied to estimate the exponent. Linear regression was performed using the "lm" function from the GLM.jl package in Julia to infer the slope (equal to the exponent of the power law) and an intercept. Non-linear regression was performed using the "curve_fit" function from the LsqFit.jl package in Julia and by passing a model of the form of Eq. (53), to infer the prefactor $A$ and exponent $n$.

If any of these simulated data points contained a mean of 0 then this simulation batch was rejected (due to issues with linear regression in log-log space). This procedure was repeated until a total of 5,000 estimates were obtained, of which the 10th, 25th, 50th, 75th, and 90th percentiles are shown in the boxplots in Fig. S2-S5. Fig. S2 and Fig. S4 show the results of linear regression (blue) to estimate the exponent for the model configurations $N = 5, j = 4$; and $N = 5, j = 1$, respectively. Fig. S3 and Fig. S5 show the results of non-linear regression (yellow) to estimate the exponent for the model configurations $N = 5, j = 4$; and $N = 5, j = 1$, respectively. In each of these figures, each row represents parameter sets whose mean at the final time point was approximately: 1 (A-E), 5 (F-J), 10 (K-O), and 25 (P-T) for Figs. S2 and S3; and 1 (A-E), 5 (F-J), 8 (K-O), and 12 (P-T) for Figs. S4 and S5.

##### 6.3 Figs. S6-S9 - Dependence of the short-time exponent estimate on mRNA capture probability

To investigate the effects of mRNA capture success on the estimation of the short-time exponent, a similar procedure as that detailed in SI Note 6. 2 was performed. The mean mRNA count was simulated for the same 20 parameter sets but the finite sample size was fixed to 1000 cells for all simulations. The binomial capture success probability, $p$, was varied to represent the non-perfect capture of mRNAs by single-cell mRNA measurements e.g. smFISH or scRNA-seq ($p = 0.2$, 0.4, 0.6, 0.8, 1.0). Given an integer mRNA count $n$ for a cell at a time point from the SSA, we generated the corresponding observed mRNA count by drawing a binomial random number with success probability $p$ and a number of trials equal to $n$ [12]. Repeated application of the binomial sampling procedure, followed by regression was used to obtain a total of 5000 estimates of the exponent for each parameter set and value of $p$. This were then used to compute the sampling distributions summarised by boxplots in Figs. S6-S9. Note that these show the 10th, 25th, 50th, 75th, and 90th percentiles.

Fig. S6 and Fig. S8 show the results of linear regression (blue) to estimate the exponent for the model configurations $N = 5, j = 4$; and $N = 5, j = 1$ respectively. Fig. S7 and Fig. S9 show the results of non-linear regression (yellow) to estimate the exponent for the model configurations $N = 5, j = 4$; and $N = 5, j = 1$ respectively. In each of these figures, each row represents parameter sets whose mean at the final time point was approximately: 1 (A-E), 5 (F-J), 10 (K-O), and 25 (P-T) for Figs. S6 and S7; and 1 (A-E), 5 (F-J), 8 (K-O), and 12 (P-T) for Figs. S8 and S9.

##### 6.4 Figs. S10-S11 - Additional results from fitting models to the yeast data

As mentioned briefly in SI Note 5.4, to determine the optimal number of time points to use in the regression we choose the maximum number of time points that minimises the RSS between the data and the model. The RSS is defined as [13]:

$$S = \sum_{i=1}^{T} r_i^2, \tag{92}$$

where $T$ is the total number of time points considered and the residual for time point $i$ is $r_i = m_i - f(t_i; A, n, t_0, m_0)$, the difference between the observed $m_i$ mean mRNA count at time $t_i$ and the corresponding regressed value given by the model.

Fig. S10 shows each induction curve (above) from the yeast data and the corresponding RSS vs number of data points plots (below). The optimal number of data points has been coloured for each curve and can visually be seen as the value after a steep decrease in the RSS (from the right). Since the nuclear mRNA curves were fit after the delay was inferred from the cytoplasmic mRNA curves, the number of data points was first optimised for the corresponding cytoplasmic curve, and subsequently optimised for the nuclear curve.

Fig. S11 shows the sampling distributions of the model parameters not displayed in Fig. 5 in the main text. These distributions were computed using 10,000 independent bootstrapped samples of the nuclear and cytoplasmic mRNA induction curves. Fig. S11A-D show the difference between the estimated nuclear and cytoplasmic exponents. Fig. S11E-H show the distributions of the fixed time delay before transcription initiation. Fig. S11I-L and Q-T show the distributions of the nuclear and cytoplasmic mRNA prefactors respectively. Fig. S11M-P and U-X show the distributions of the mean nuclear and cytoplasmic mRNA counts during the delay respectively.

##### 6.5 Fig. S12 - Comparing the results of fitting models with and without a delay before transcription initiation

If the time delay to the beginning of transcription initiation is unknown, then it is important to infer this delay, else the exponent (and subsequent number of gene states) may be overestimated. Fig. S12 shows the results of comparing the fits from a model with no fixed time delay (red) to one with a fixed time delay (blue, same as in Fig. 5 in the main text) to the CTL1 (0.2 mol NaCl) yeast data. The model with no fixed delay takes the form of Eq. (53), where only the prefactor and exponent were inferred. The same bootstrapping procedure as detailed in SI Note 5.4 was used to generate the sampling distributions for the models with delay and with no fixed delay.

Fig. S12A and B show the nuclear and cytoplasmic mean mRNA induction curves respectively with the regression given by the two different models. In each panel the fits from the two different models are indistinguishable from each other, despite the clear difference in the exponents. Fig. S12C and D show the sampling distributions of the nuclear and cytoplasmic exponents between the models, with clear differences between them. Fig. S12E shows the sampling distribution of the difference between the nuclear exponents estimated from the model with fixed delay and the model with no fixed delay. Fig. S12F shows the same sampling distribution but instead for the difference between the cytoplasmic exponents. The medians of both of these distributions are $> 1$, implying that at least 1 additional first-order reaction would be inferred if no fixed delay was fitted. This emphasises the importance of estimating a fixed delay before the beginning of transcription initiation to ensure accurate inferences of the exponent and subsequent number of gene states.

##### 6.6 Fig. S13 - Results from fitting models to mouse data

Data was extracted using WebPlotDigitizer [14] to obtain the pre-mRNA and mRNA counts and times from Fig. S2 of Ref. [3]. The Icam1 mRNA curve was taken from Fig. S2A and the Cxcl1 premRNA and mRNA curves were taken from Fig. S2B.

Since this data is not single cell, it was not possible to repeat the uncertainty analysis as done on the yeast data *on* this data. Instead local estimates of the uncertainty were calculated and the standard errors in each model parameter estimate reported here represent the scatter of the data about the best fit line. Additionally since each experimental replica was not available, the plots do not display error bars representing the SEM between the different experimental replicas.

The same model as used for the yeast data (see Fig. 5 in the main text, and SI Note 5.4) was fitted to these three curves to infer the prefactor, exponent, delay, and constant mean RNA count during the delay. The parameter estimates and their uncertainties can be found in Table S7.

If we denote our model parameters as the vector $\boldsymbol{\beta}$ (which has length say $U$), then the minimum RSS (see Eq. (92)) occurs when the gradient is 0, i.e. when

$$\frac{\partial S}{\partial \beta_j} = 2\sum_i r_i \frac{\partial r_i}{\partial \beta_j} = 0, \tag{93}$$

for each model parameter $j$. The Jacobian matrix, $\boldsymbol{J}$, of the linearised model [13] is given by:

$$J_{ij} = -\frac{\partial r_i}{\beta_j}. \tag{94}$$

Evaluating the Jacobian matrix at the best fit point (using the parameters that minimise the RSS, denoted by $\hat{\boldsymbol{\beta}}$) we can estimate the parameter variance-covariance matrix

$$\mathrm{Cov}(\hat{\boldsymbol{\beta}}) = \left(\boldsymbol{J}^{\boldsymbol{T}} \cdot \boldsymbol{J}\right)^{-1} \cdot \hat{\sigma}^2, \tag{95}$$

where the scalar $\hat{\sigma}^2$ can be estimated as the RSS divided by the degrees of freedom $(T - U)$ where $T$ is the total number of data points. Taking the square root of the diagonals of the estimate of the parameter variance-covariance matrix evaluated at the optimal parameter set yields estimates of the standard errors in each model parameter, i.e. measurements of local uncertainty.

##### 6.7 Fig. S14 - Comparing steady-state mRNA count distributions with equal $k_i$ rates

To compare steady-state distributions of the $N = 3, 4, 5$ state models with equal gene state switching rates, $k_i$ for $i = 1, \dots, N-1$, we chose rate parameters such that: $k_i = (N-1)a$, where $a = 0.375, 1.5, 6.0$. The remaining rate parameters were fixed at $k_N = 1.5$, $\rho = 100$, and $d = 1$. This forced the fraction of cells in the active state, denoted $f_{\mathrm{ON}}$, to be 0.2, 0.5, and 0.8, depending on the value of $a$. Note that $f_{\mathrm{ON}} = \sigma_u/(\sigma_u + \sigma_b)$ where $\sigma_u$ and $\sigma_b$ are the expressions for the effective gene inactivation and activation activation rates described in SI Note 5.1. See Table S8 for a summary of the rate parameters. Fig. S14 shows these distributions (blue) and their effective telegraph distributions (red) for the $N = 3$ (A, D, G), $N = 4$ (B, E, H), and $N = 5$ (C, F, I) state models. In all cases, the $N$-state model steady-state mRNA count distribution is well approximated by that of the effective telegraph model, both qualitatively and quantitatively. For each fixed $f_{\mathrm{ON}}$, the Wasserstein distance (WD) between the $N$-state and effective telegraph distribution increases as $N$ increases, but is still well approximated by the effective telegraph. Interestingly, for each fixed $f_{\mathrm{ON}}$ and all $N$, the distributions look very similar, indicating that it is difficult to infer the number of rate limiting steps in steady-state.

##### 6.8 Figs. S15-S16 - Effect of mixed gene state initial conditions on short-time exponent estimation

Fig. S15 shows the short-time behaviour of the mean mRNA count for mixed gene state initial conditions of a 5-state model for one particular parameter set (see caption for the parameter values). Here, for simplicity, we consider that the gene initially is in state $G_1$ with probability $P_{G_1}(0) = 1 - x$ and in state $G_3$ with probability $P_{G_3}(0) = x$ where $x$ is a fraction. Fig. S15A and B show the mean mRNA count curves for different values of $x$, computed by direct numerical integration of the first moment equation of the CME on log-log and linear scales, respectively.

In Fig. S15A the $x = 0.0$ (purple) line has a slope of 5 over very short times, since all cells begin in state $G_1$ and four first-order reactions are needed for gene activation and a fifth for transcription. Similarly the $x = 1.0$ (yellow) line has a slope of 3 over very short times (all cells begin in state $G_3$ so two first-order reactions are needed for gene activation and another for transcription). For $0 < x < 1$, over very short times, we see each line parallel to the $x = 1.0$ line, indicating that the slope (and thus exponent) is also 3 for all of these initial conditions (over very short times). It follows by the results of SI Note 2.2, that this is because for all initial conditions except the case $x = 0$, there is a finite non-zero probability of initially being in state $G_3$. For longer and more experimentally reasonable times (see dots for sample data) deviations from an exponent equal to 3 appear because of the non-zero probability of initially being in state $G_1$. While Eq. (45) cannot analytically be reduced to a power law, nevertheless we find that it can very well be approximated by one in many cases — we confirm this by showing that fitting a power law to the five sampled measurements using linear or non-linear regression leads to $R^2 > 0.99$ for all values of $x$ (Table S9). The corresponding estimated exponent lies between 2.39 and 4.19 (using linear regression), all below $N = 5$. Note that some values of the estimated exponent are also consistent with values expected from a Dirac delta initial state distribution with say $j = 2$ (all cells in state $G_2$), despite the initial gene state being *only* either $G_1$ or $G_3$; hence generally it is difficult to tell apart a mixed initial gene state distribution from a Dirac delta initial state distribution. However because the exponent is greater than 2 for all $x$, it means that for these particular mixed initial conditions, we can distinguish the 5-state model from the telegraph model. In Fig. S15C we show that similar results are obtained when we account for the finite sample size by fitting a single power law to simulated data from 1000 cells using the SSA.

We also repeated the numerical experiment using the initial condition that the gene initially is in state $G_1$ with probability $P_{G_1}(0) = 1 - x$ and in state $G_4$ with probability $P_{G_4}(0) = x$. The best fit exponents (Table S10) are found to be larger than 2 only when $x \leq 0.05$; this means that in this case, it is very difficult to tell apart the $N$-state model from a telegraph model because only a small probability of being in $G_4$ is enough to shift the exponent to a value close to or below 2. Increasing the number of sampled time points makes no difference to this result.

To understand more generally how difficult it is to distinguish the telegraph model from the 5-state model with initial gene state distributions with a non-zero probability of being in state $G_4$, we performed a sweep across parameter space (similar to that done for Fig. 4 in the main text, see SI Note 5.3 for the procedure) whilst also varying the initial gene state distribution (illustrated in Fig. S16B-G) in 2 different ways: (i) Approximately uniform distributions (Fig. S16G) were obtained by choosing four uniformly distributed random numbers in the interval $(0, 1]$, normalised by their sum to determine the initial probability in each one of the four inactive states ($G_1$, ..., $G_4$); (ii) Distributions concentrated in one state (Fig. S16C and F) were obtained by drawing three random numbers from a log-uniform distribution on the interval $[10^{-5}, 10^{-1}]$ to be the probabilities of starting in the remaining three states and then subtracted their sum from 1 to give the probability of starting in the most probable state, provided this was $> 0.9$. Note that in all cases the probability of a cell initially being in state $G_5$ (the only transcriptionally active state) was fixed to be 0 and the initial mean mRNA count was also fixed to be 0. Using these sampling procedures, we generated 33 different initial distributions of type (i), 33 distributions of type (ii) mostly in the $G_1$ state and 33 distributions of type (ii) mostly in the $G_4$ state. These were combined with the three remaining cases (Dirac delta in state $G_1$, Dirac delta in state $G_4$, and exact uniform illustrated in Fig. S16B, E, D respectively) resulting in 102 initial conditions for each of the 100 different parameter sets used in Fig. S16. For each of these, we simulated the mean mRNA count (by direct numerical integration of the first moment equations of the CME) and performed a linear regression in log-log space to extract the exponent of the power law. In Fig. S16A we show a scatter plot of the estimated exponent against the standard deviation of the initial gene state distribution $\sigma_{u_0}$:

$$\sigma_{u_0} = \sqrt{\sum_{i=1}^{N} i^2 P_{G_i}(0) - \left(\sum_{i=1}^{N} i P_{G_i}(0)\right)^2}, \tag{96}$$

which is a measure of how concentrated the initial distribution is in a single gene state.

The main conclusions from the analysis shown in Fig. S16 are in agreement with the previous results in Table S9, Table S10 and Fig. S15: (i) a single power law can be very well fit (95% of regressions had an $R^2 \geq 0.99$ in Fig. S16H); (ii) in all cases, the exponent is below the number of gene states $N = 5$; (iii) distinguishing the 5-state model from the telegraph model (exponent $> 2$) is mostly possible when the initial distribution is concentrated in the $G_1$ state and the standard deviation was below $\approx 0.5$. A small fraction of the approximately uniform initial distributions also gave rise to exponents $> 2$; for these parameter sets, the median initial probabilities of being in states $G_1$, $G_2$, $G_3$ and $G_4$ are 0.26, 0.30, 0.37 and 0.043, respectively. Hence generally a Dirac-delta-like initial gene state distribution is not necessary to distinguish the telegraph from higher state models so long as the probability of initially being in state $G_4$ is small. The underlying reason for this limitation stems from the result in SI Note 2.2 which says that if $P_{G_{N-1}}(0) \neq 0$ then the leading order term in Eq. (45) is quadratic in time ($t^2$) which is the same as from a telegraph model. Hence, generally we expect the method to be particularly useful when the repressor used to suppress gene activity (prior to induction) forces the system into a deep inactive state, i.e. $G_1, ..., G_{N-2}$ (see for example Ref. [15]).

### SUPPLEMENTARY TABLES

| Parameter | Shape | | | |
|---|---|---|---|---|
| | I | II | III | IV |
| $k_1$ | 29 | 7.2 | 1.2 | 2.94 |
| $k_2$ | 16.3 | 17.3 | 0.9 | 4.21 |
| $k_3$ | 37.5 | 12.1 | 1.1 | 2.6 |
| $k_4$ | 36.7 | 32 | 1.4 | 1.7 |
| $k_5$ | 0.1 | 5.4 | 0.25 | 0.58 |
| $\rho$ | 35 | 81 | 40 | 80 |
| $d$ | 1 | 1 | 1 | 1 |
| Wasserstein distance | 0.00588 | 0.235 | 0.946 | 1.44 |

TABLE. S1. Rate parameter values used for steady-state mRNA count distribution shapes I-IV in Fig. 2D-G.

| Distribution Shape | $N$-state model | Percentile | | | | | Number of samples |
|---|---|---|---|---|---|---|---|
| | | 10th | 25th (LQ) | 50th (median) | 75th (UQ) | 90th | |
| I | 3 | 0.00161 | 0.00442 | 0.0106 | 0.0246 | 0.0471 | 151 |
| II | | 0.00196 | 0.00955 | 0.0520 | 0.161 | 0.400 | 1053 |
| III | | 0.00179 | 0.0106 | 0.0792 | 0.308 | 0.716 | 183 |
| IV | | 0.270 | 0.500 | 1.02 | 1.86 | 2.65 | 243 |
| I | 4 | 0.00670 | 0.0115 | 0.0280 | 0.0634 | 0.108 | 313 |
| II | | 0.0159 | 0.0552 | 0.172 | 0.413 | 0.834 | 6681 |
| III | | 0.0212 | 0.0948 | 0.302 | 0.783 | 1.25 | 1313 |
| IV | | 0.330 | 0.571 | 1.09 | 1.68 | 2.24 | 509 |
| I | 5 | 0.0124 | 0.0230 | 0.0508 | 0.0862 | 0.129 | 963 |
| II | | 0.0409 | 0.110 | 0.262 | 0.614 | 1.13 | 30758 |
| III | | 0.0753 | 0.207 | 0.471 | 0.952 | 1.60 | 7023 |
| IV | | 0.427 | 0.692 | 1.25 | 1.96 | 2.66 | 1464 |

TABLE. S2. Statistics summarising the WD distances computed between the steady-state distributions of the effective telegraph and the $N = 3, 4, 5$ state models using randomly chosen rate parameter values covering the physiologically relevant parameter space. See Fig. 2H for the box plots displaying these results.

| Parameter | Value | | | |
|---|---|---|---|---|
| | Panels A, B, C | Panel D | Panel F ($t^3$) | Panel F ($t^4$) |
| $k_1$ | 18.5 | 18.5 | 18.5 | 20.5 |
| $k_2$ | 12.5 | 12.5 | 22.5 | 35.5 |
| $k_3$ | 14.5 | 14.5 | 14.5 | 44.5 |
| $k_4$ | 13.5 | 13.5 | 17.5 | 28.0 |
| $k_5$ | 1.7 | 1.7 | 4.0 | 0.5 |
| $\rho$ | 850 | 850 | 1000 | 1600 |
| $d$ | 1.0 | 1.0 | 1.0 | 1.0 |
| $\gamma$ | - | 100 | - | - |
| $d_p$ | - | 0.2 | - | - |
| $k_{2,5}$ | - | - | 9.0 | - |
| $k_{2,1}$ | - | - | - | 0.5 |

TABLE. S3. Rate parameter values used in Fig. 3A-D, and F.

| Rate parameters | Gamma distribution | | | | | | |
|---|---|---|---|---|---|---|---|
| | Mean value | $t^2$ | | $t^4$ | | $t^5$ | |
| | | Shape | Scale | Shape | Scale | Shape | Scale |
| $k_1$ | 18.5 | 0.25 | 74 | 4 | 4.625 | 1 | 18.5 |
| $k_2$ | 12.5 | 0.25 | 50 | 4 | 3.125 | 1 | 12.5 |
| $k_3$ | 14.5 | 0.25 | 58 | 4 | 3.625 | 1 | 14.5 |
| $k_4$ | 13.5 | 0.25 | 54 | 4 | 3.375 | 1 | 13.5 |
| $k_5$ | 1.7 | 0.25 | 6.8 | 4 | 0.425 | 1 | 1.7 |
| $\rho$ | 850 | 0.25 | 3400 | 4 | 212.5 | 1 | 850 |
| $d$ | 1.0 | - | - | - | - | - | - |

TABLE. S4. Rate parameter and gamma distribution parameter values used in Fig. 3E.

| Parameter | Value | | | | | | | | | | | |
|---|---|---|---|---|---|---|---|---|---|---|---|---|
| $k_1$ | 2.6 | 3.8 | 24.5 | 1.35 | 1.72 | 4.3 | 19.7 | 0.52 | 0.16 | 1.67 | 3.28 | 2.5 |
| $k_2$ | 27 | 2.2 | 27.2 | 5.6 | 4.7 | 0.12 | 0.95 | 38.2 | 6 | 2.44 | 3.25 | 2.5 |
| $k_3$ | 19.6 | 9.1 | 25.7 | 5.1 | 5.2 | 1.45 | 29.4 | 7.97 | 2.41 | 1.97 | 2.36 | 2.5 |
| $k_4$ | 14 | 7.1 | 27.5 | 25 | 1.4 | 1.36 | 0.12 | 6.56 | 0.54 | 1.24 | 3.66 | 2.5 |
| $k_5$ | 101 | 0.13 | 0.27 | 0.82 | 0.15 | 15.5 | 0.11 | 0.1 | 0.14 | 0.37 | 0.54 | 0.625 |
| $\rho$ | 135 | 21 | 195 | 192 | 170 | 343 | 124 | 77 | 8 | 155 | 49 | 100 |
| $d$ | 1 | 1 | 1 | 1 | 1 | 1 | 1 | 1 | 1 | 1 | 1 | 1 |
| WD | 0.00911 | 0.0946 | 0.276 | 1.71 | 2.21 | 0.0787 | 0.402 | 0.129 | 0.0632 | 3.53 | 0.808 | 1.97 |
| Shape | I | | | II | | | III | | | IV | | |
| Fig. S1 Panel | A | E | I | B | F | J | C | G | K | D | H | L |

TABLE. S5. Rate parameter values of the $N$-state model ($N = 5$) used for the steady-state mRNA count distributions in Fig. S1.

| Time (mins) | Number of cells | |
|---|---|---|
| | Exp 1 (0.2 mol NaCl) | Exp 2 (0.4 mol NaCl) |
| 0 | 3958 | 2001 |
| 1 | 2214 | 4771 |
| 2 | 1807 | 2545 |
| 4 | 1962 | 1355 |
| 6 | 1627 | 2509 |
| 8 | 1747 | 1476 |
| 10 | 1254 | 3370 |
| 15 | 1397 | 4813 |

TABLE. S6. The number of cells measured at each time point after combining experimental replicas for the yeast data [4] used in Fig. 5 and Figs. S10-S12.

| Cell type | Gene | RNA species | Estimate ± standard error | | | | Fig. S13 Panel |
|---|---|---|---|---|---|---|---|
| | | | Exponent | Delay (mins) | Prefactor | $m_0$ | |
| Macrophage | Cxcl1 | premRNA | 1.682 ± 0.204 | 4.321 ± 1.798 | 2.426 ± 0.685 | $2.21 \times 10^{-15} \pm 1.18$ | A |
| Macrophage | Cxcl1 | mRNA | 2.506 ± 0.605 | 4.321 ± 1.798 | 1.322 ± 1.840 | 3.936 ± 3.981 | B |
| Fibroblast | Icam1 | mRNA | 2.749 ± 0.356 | 13.41 ± 4.20 | 0.00448 ± 0.00583 | $3.69 \times 10^{-16} \pm 0.551$ | C |

TABLE. S7. Parameter estimates and their standard errors from the results of fitting the model in Eq. (87) to mouse data [3] used in Fig. S13.

| Parameter | Value | | | | | | | | |
|---|---|---|---|---|---|---|---|---|---|
| $a$ | 0.375 | 1.5 | 6.0 | 0.375 | 1.5 | 6.0 | 0.375 | 1.5 | 6.0 |
| $k_i$ | 0.75 | 3.0 | 12.0 | 1.125 | 4.5 | 18.0 | 1.5 | 6.0 | 24.0 |
| $k_N$ | 1.5 | 1.5 | 1.5 | 1.5 | 1.5 | 1.5 | 1.5 | 1.5 | 1.5 |
| $\rho$ | 100 | 100 | 100 | 100 | 100 | 100 | 100 | 100 | 100 |
| $d$ | 1 | 1 | 1 | 1 | 1 | 1 | 1 | 1 | 1 |
| $f_{\mathrm{ON}}$ | 0.2 | 0.5 | 0.8 | 0.2 | 0.5 | 0.8 | 0.2 | 0.5 | 0.8 |
| $N$ | 3 | 3 | 3 | 4 | 4 | 4 | 5 | 5 | 5 |
| WD | 0.809 | 1.175 | 1.389 | 0.628 | 0.888 | 1.031 | 0.189 | 0.253 | 0.285 |
| Fig. S14 Panel | A | D | G | B | E | H | C | F | I |

TABLE. S8. Table of $N$-state Rate parameters used in Fig. S14.

| Initial condition | | Linear regression (Fig. S15A) | | Non-linear regression (Fig. S15B) | |
|---|---|---|---|---|---|
| $P_{G_1}(0)$ | $P_{G_3}(0)$ | $R^2$ | Estimated exponent (slope) | $R^2$ | Estimated exponent |
| 0.0 | 1.0 | 0.997 | 2.39 | 0.999 | 2.12 |
| 0.5 | 0.5 | 0.999 | 2.50 | 1.00 | 2.30 |
| 0.8 | 0.2 | 1.00 | 2.73 | 1.00 | 2.66 |
| 0.9 | 0.1 | 1.00 | 2.97 | 1.00 | 2.97 |
| 0.95 | 0.05 | 1.00 | 3.25 | 1.00 | 3.24 |
| 0.98 | 0.02 | 1.00 | 3.61 | 1.00 | 3.47 |
| 0.99 | 0.01 | 1.00 | 3.82 | 1.00 | 3.56 |
| 0.995 | 0.005 | 0.999 | 3.98 | 1.00 | 3.61 |
| 0.998 | 0.002 | 0.999 | 4.10 | 1.00 | 3.64 |
| 0.999 | 0.001 | 0.999 | 4.14 | 1.00 | 3.65 |
| 1.0 | 0.0 | 0.998 | 4.19 | 1.00 | 3.66 |

TABLE. S9. Results of linear and non-linear regression applied to five sampled mean mRNA count measurements computed from direct numerical integration of the first moment equations of the CME for various mixed gene state initial conditions. The probability that a gene is initially in state $G_1$ is $P_{G_1}(0) = 1 - x$ and the probability that it is in state $G_3$ is $P_{G_3}(0) = x$. To investigate different initial conditions $x$ is varied between 0.0 and 1.0. The value of $R^2$ (3 s.f.) and the estimated slope (linear regression) or exponent (non-linear regression) is given for each initial condition. The following rate parameters of the $N = 5$-state model were used: $k_1 = 18.6, k_2 = 18.3, k_3 = 13.0, k_4 = 29.7, k_5 = 1.3, \rho = 880, d = 1.0$. The mean mRNA count curves are seen in Fig. S15A-B. See SI Note 6.8 for further details.

| Initial condition | | Linear regression | | Non-linear regression | |
|---|---|---|---|---|---|
| $P_{G_1}(0)$ | $P_{G_4}(0)$ | $R^2$ | Estimated exponent (slope) | $R^2$ | Estimated exponent |
| 0.0 | 1.0 | 0.997 | 1.50 | 0.998 | 1.36 |
| 0.5 | 0.5 | 0.998 | 1.55 | 0.999 | 1.45 |
| 0.8 | 0.2 | 1.00 | 1.68 | 1.00 | 1.70 |
| 0.9 | 0.1 | 0.998 | 1.85 | 0.999 | 2.03 |
| 0.95 | 0.05 | 0.995 | 2.10 | 0.999 | 2.46 |
| 0.98 | 0.02 | 0.993 | 2.53 | 1.00 | 3.01 |
| 0.99 | 0.01 | 0.994 | 2.88 | 1.00 | 3.29 |
| 0.995 | 0.005 | 0.997 | 3.23 | 1.00 | 3.47 |
| 0.998 | 0.002 | 0.999 | 3.62 | 1.00 | 3.58 |
| 0.999 | 0.001 | 1.00 | 3.84 | 1.00 | 3.62 |
| 1.0 | 0.0 | 0.998 | 4.19 | 1.00 | 3.66 |

TABLE. S10. Results of linear and non-linear regression applied to five sampled mean mRNA count measurements computed from direct numerical integration of the first moment equations of the CME for various mixed gene state initial conditions. The probability that a gene is initially in state $G_1$ is $P_{G_1}(0) = 1 - x$ and the probability that it is in state $G_4$ is $P_{G_4}(0) = x$. To investigate different initial conditions $x$ is varied between 0.0 and 1.0. The value of $R^2$ (3 s.f.) and the estimated slope (linear regression) or exponent (non-linear regression) is given for each initial condition. The following rate parameters of the $N = 5$-state model were used: $k_1 = 18.6, k_2 = 18.3, k_3 = 13.0, k_4 = 29.7, k_5 = 1.3, \rho = 880, d = 1.0$. See SI Note 6.8 for further details.

### SUPPLEMENTARY FIGURES

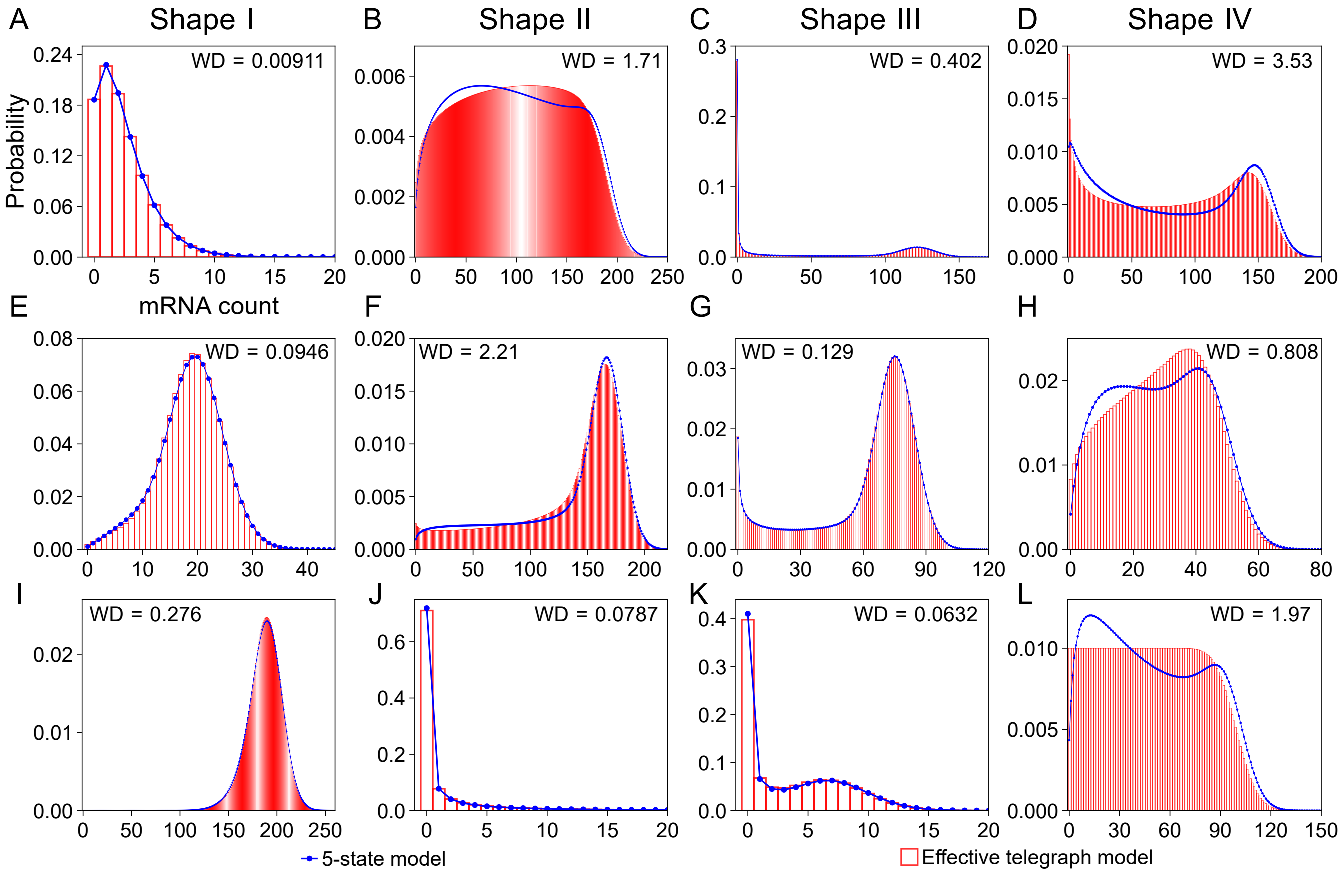

Fig. S1. Steady-state mRNA count distributions from the $N$-state model with $N = 5$ come in a variety of shapes, many of which can be well approximated by an effective telegraph model. Each panel displays the Wasserstein distance (WD) between the 5-state model (solid blue lines) and the effective telegraph model (histograms) distributions for three cases of each shape class. The parameters of the effective telegraph model are determined using Eq. (2) in the main text. Histogram bar widths represent a single mRNA value. (A, E, I) Shape I - unimodal with Fano factor $< 2$; (B, F, J) Shape II - unimodal with Fano factor $\geq 2$; (C, G, K) Shape III - bimodal with one mode at zero value and one mode at a non-zero value; (D, H, L) Shape IV - bimodal with both modes at non-zero values. See SI Note 6.1 for further details and Table S5 for the rate parameter values for each distribution.

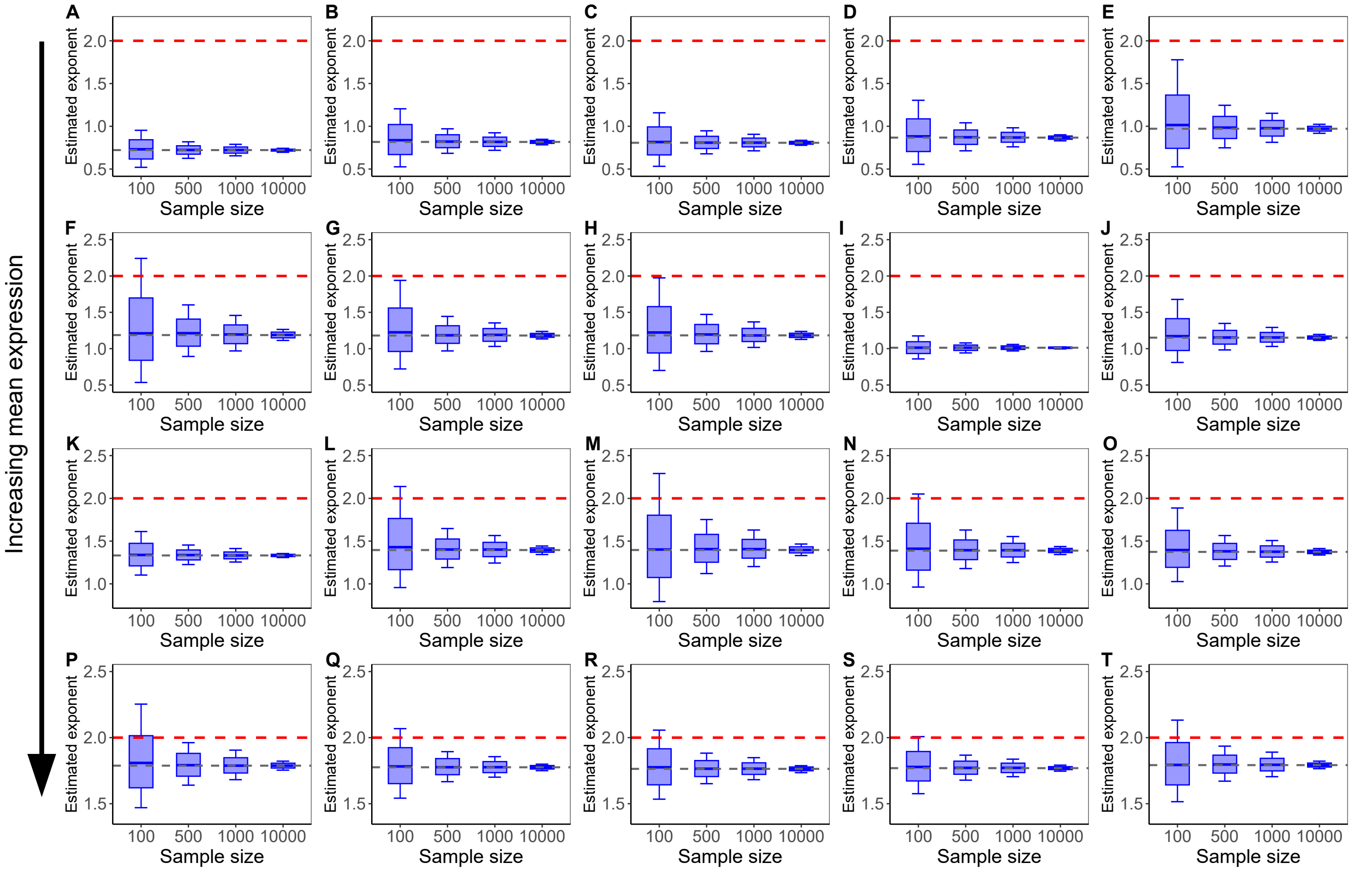

Fig. S2. Increasing the number of cells sampled improves the estimate of the exponent from linear regression on the mean mRNA count following induction. Panels A-T show the estimated exponent using linear regression from 20 different parameters sets using 5000 independent samples of the mean computed over different sample sizes (100, 500, 1000, 10000), generated using the SSA. Boxplots show the median, interquartile range (IQR), and whiskers extended to the 10th and 90th percentiles. The grey dashed horizontal line shows the infinite sample size exponent and the red line shows the theoretical short time exponent $(N - j + 1)$. In this case $N = 5, j = 4$. Each row represents parameter sets whose mean at the final sample was approximately 1 (A-E), 5 (F-J), 10 (K-O), and 25 (P-T). Parameter sets were chosen from those randomly generated for Fig. 4. See SI Note 6. 2 for details on the sampling procedure.

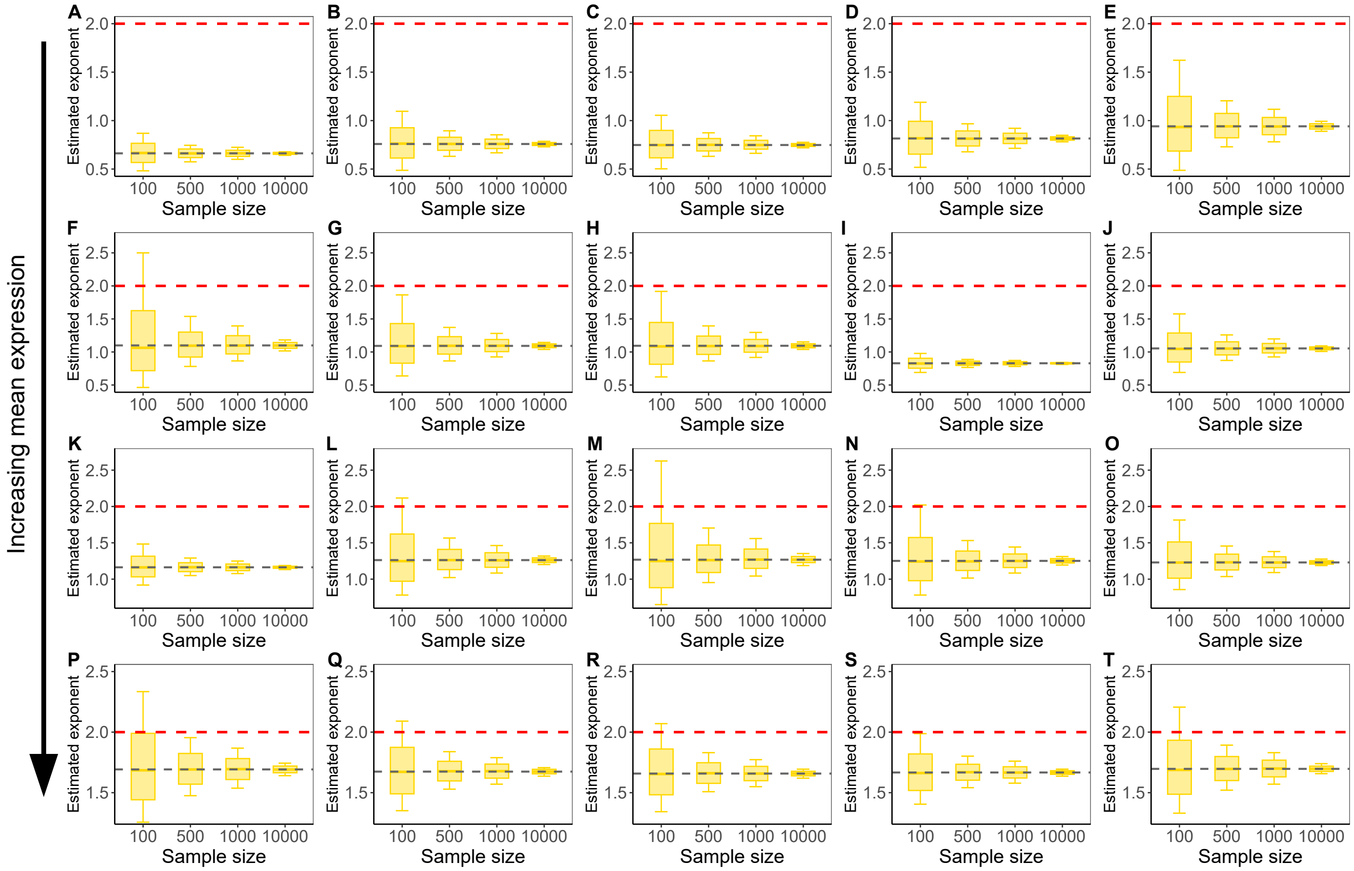

Fig. S3. Increasing the number of cells sampled improves the estimate of the exponent from non-linear regression on the mean mRNA count following induction. Panels A-T show the estimated exponent using non-linear regression from 20 different parameters sets using 5000 independent samples of the mean computed over different sample sizes (100, 500, 1000, 10000), generated using the SSA. Boxplots show the median, IQR, and whiskers extended to the 10th and 90th percentiles. The grey dashed horizontal line shows the infinite sample size exponent, and the red line shows the theoretical short time exponent $(N - j + 1)$. In this case $N = 5, j = 4$. The distribution gets narrower as the sample size increases. Sample sizes of 500 or greater are recommended. Each row represents parameter sets whose mean at the final sample was approximately 1 (A-E), 5 (F-J), 10 (K-O), and 25 (P-T). Parameter sets were chosen from those randomly generated for Fig. 4. See SI Note 6.2 for details on the sampling procedure..

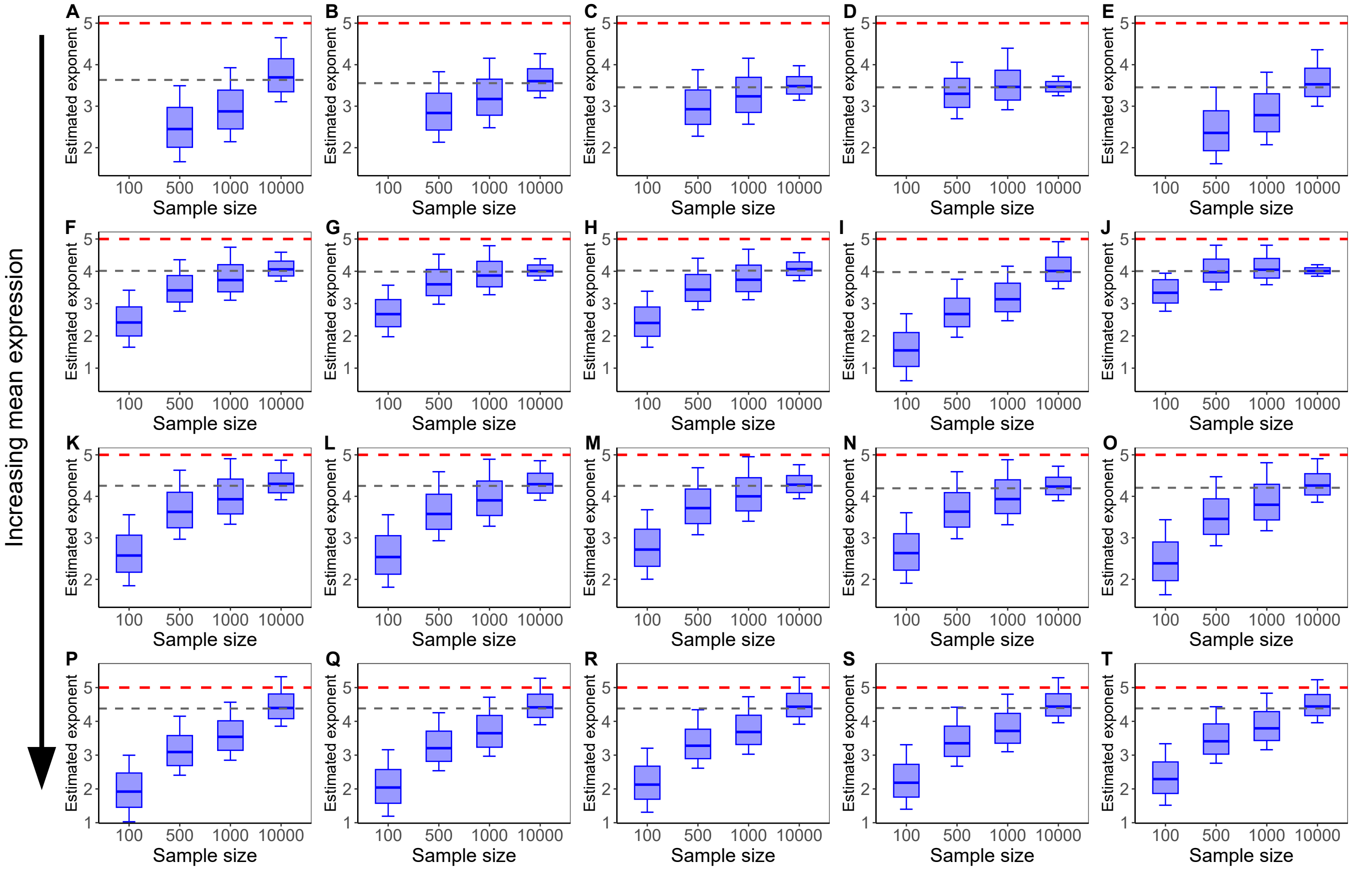

Fig. S4. Increasing the number of cells sampled improves the estimate of the exponent from linear regression on the mean mRNA count following induction. Panels A-T show the estimated exponent using linear regression from 20 different parameters sets using 5000 independent samples of the mean computed over different sample sizes (100, 500, 1000, 10000), generated using the SSA. Boxplots show the median, IQR, and whiskers extended to the 10th and 90th percentiles. The grey dashed horizontal line shows the infinite sample size exponent, and the red line shows the theoretical short time exponent $(N - j + 1)$. In this case $N = 5, j = 1$. The distribution of estimated exponents moves substantially closer to the infinite sample size estimate as the sample size increases. Sample sizes of 500 or greater are recommended. Each row represents parameter sets whose mean at the final sample was approximately 1 (A-E), 5 (F-J), 8 (K-O), and 12 (P-T). Parameter sets were chosen from those randomly generated for Fig. 4. See SI Note 6.2 for details on the sampling procedure.

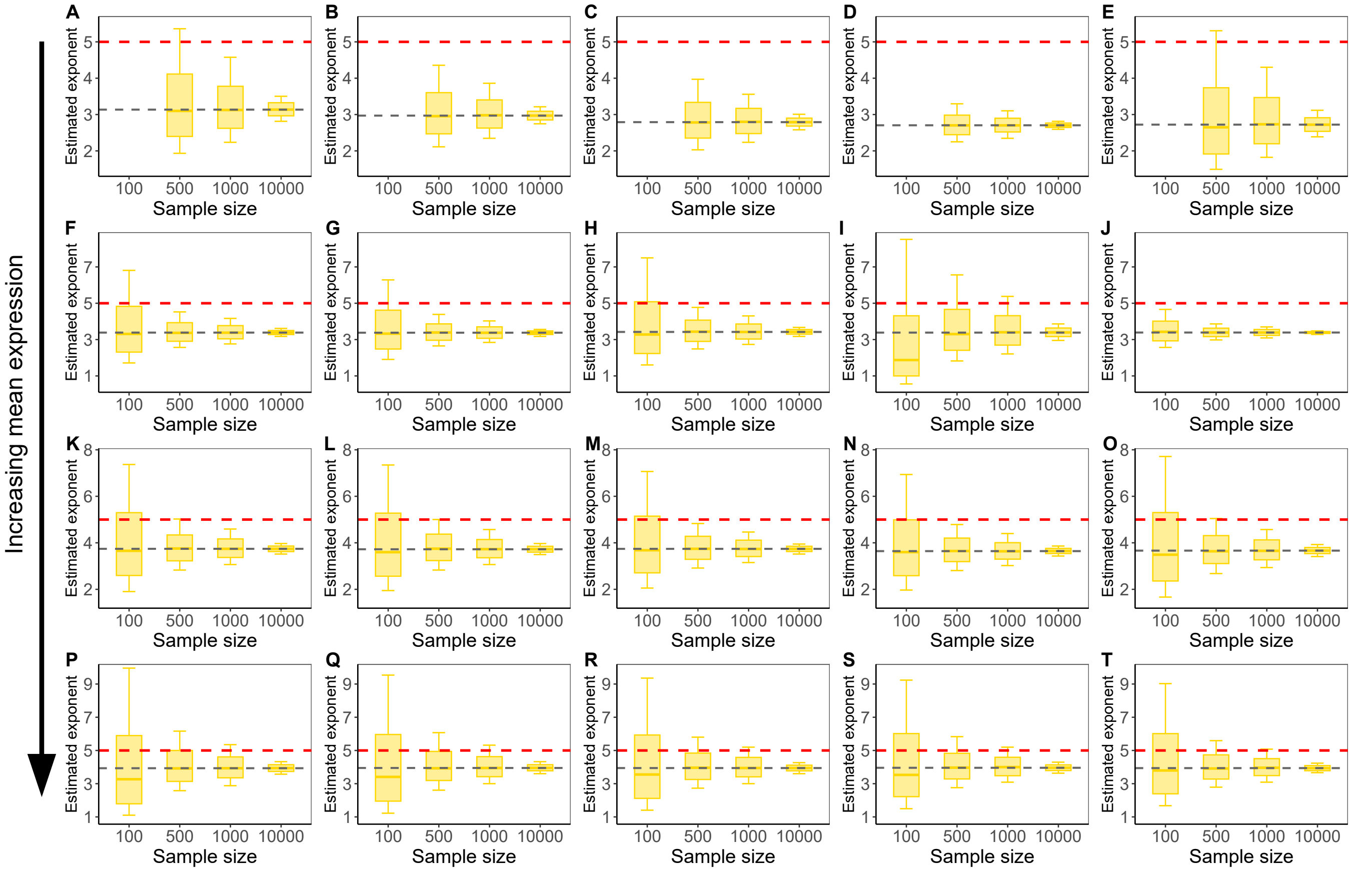

Fig. S5. Increasing the number of cells sampled improves the estimate of the exponent from non-linear regression on the mean mRNA count following induction. Panels A-T show the estimated exponent using non-linear regression from 20 different parameters sets using 5000 independent samples of the mean computed over different sample sizes (100, 500, 1000, 10000), generated using the SSA. Boxplots show the median, IQR, and whiskers extended to the 10th and 90th percentiles. The grey dashed horizontal line shows the infinite sample size exponent, and the red line shows the theoretical short time exponent $(N - j + 1)$. In this case $N = 5, j = 1$. The distribution gets narrower as the sample size increases. Sample sizes of 500 or greater are recommended. Each row represents parameter sets whose mean at the final sample was approximately 1 (A-E), 5 (F-J), 8 (K-O), and 12 (P-T). Parameter sets were chosen from those randomly generated for Fig. 4. See SI Note 6.2 for details on the sampling procedure.

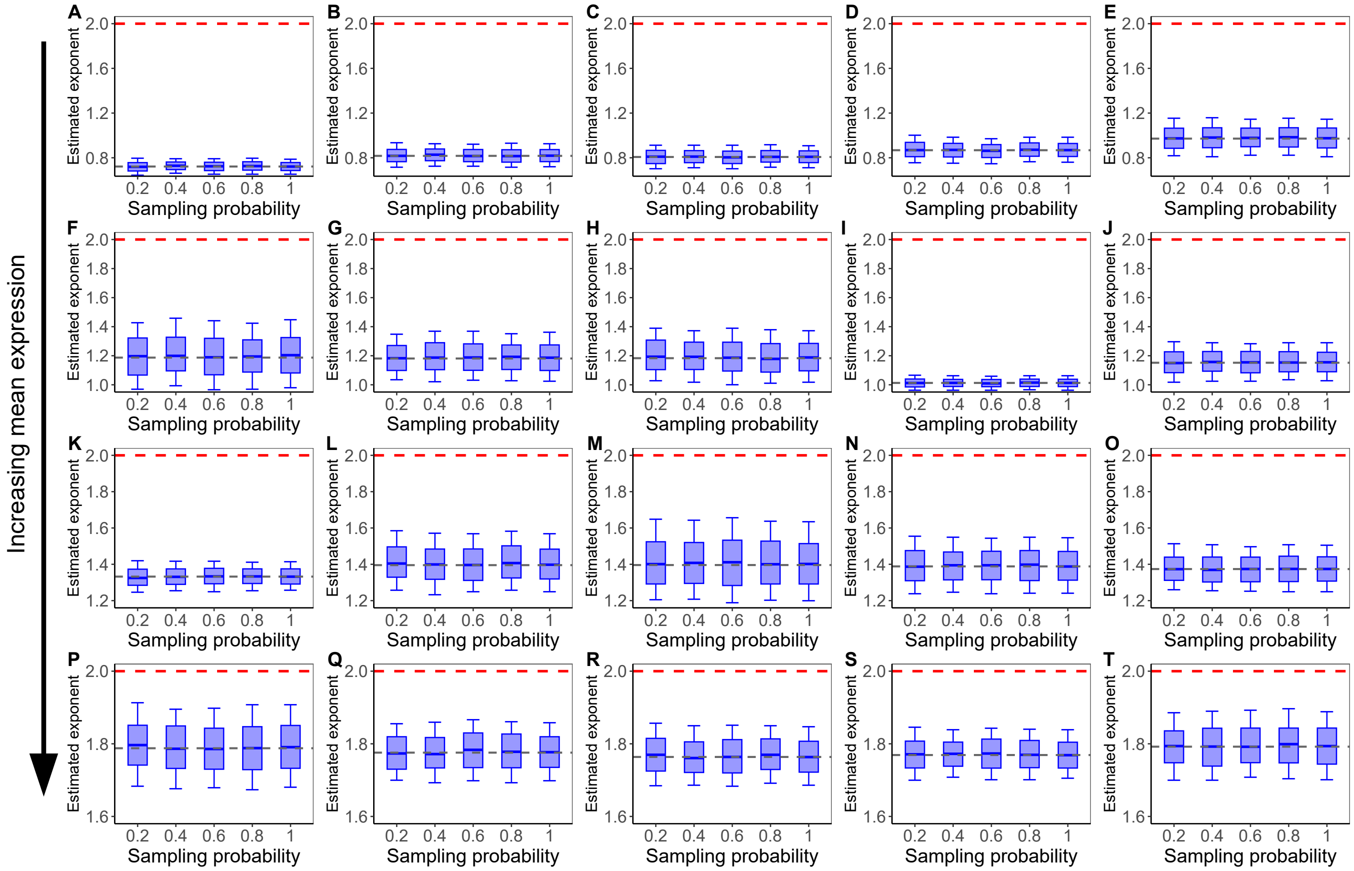

Fig. S6. Changing the mRNA capture success probability does little to affect the estimated exponent from linear regression on the mean mRNA count following induction. Panels A-T show the estimated exponent using linear regression from 20 different parameters sets using 5000 independent samples of the mean computed with sample size (number of cells) 1000, generated using the SSA. Each sample was resampled using binomial sampling procedure representing technical noise introduced by the mRNA capture procedure. The following binomial sampling probabilities of 0.2, 0.4, 0.6, 0.8, and 1.0 were used to represent various mRNA capture success rates. Boxplots show the median, IQR, and whiskers extended to the 10th and 90th percentiles. The grey dashed horizontal line shows the infinite sample size exponent, and the red line shows the theoretical short time exponent $(N - j + 1)$. In this case $N = 5, j = 4$. The distribution of estimated exponents remains the same as the sampling probability increases. Each row represents parameter sets whose mean at the final sample was approximately 1 (A-E), 5 (F-J), 10 (K-O), and 25 (P-T). Parameter sets were chosen from those randomly generated for Fig. 4. See SI Note 6.3 for details on the sampling procedure.

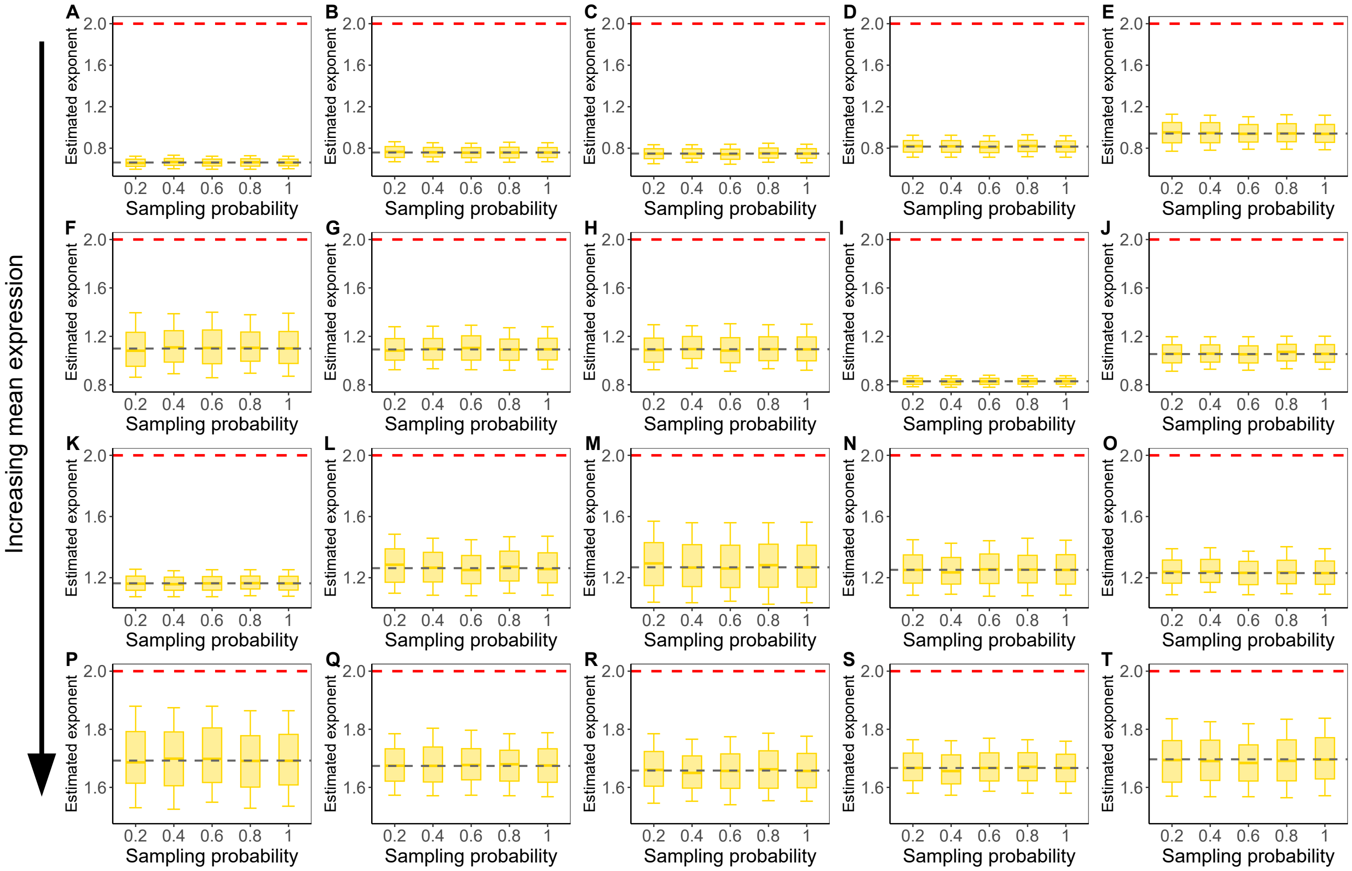

Fig. S7. Changing the mRNA capture success probability does little to affect the estimated exponent from non-linear regression on the mean mRNA count following induction. Panels A-T show the estimated exponent using linear regression from 20 different parameters sets using 5000 independent samples of the mean computed with sample size (number of cells) 1000, generated using the SSA. Each sample was resampled using binomial sampling procedure representing technical noise introduced by the mRNA capture procedure. The following binomial sampling probabilities of 0.2, 0.4, 0.6, 0.8, and 1.0 were used to represent various mRNA capture success rates. Boxplots show the median, IQR, and whiskers extended to the 10th and 90th percentiles. The grey dashed horizontal line shows the infinite sample size exponent, and the red line shows the theoretical short time exponent $(N - j + 1)$. In this case $N = 5, j = 4$. The distribution of estimated exponents remains the same as the sampling probability increases. Each row represents parameter sets whose mean at the final sample was approximately 1 (A-E), 5 (F-J), 10 (K-O), and 25 (P-T). Parameter sets were chosen from those randomly generated for Fig. 4. See SI Note 6.3 for details on the sampling procedure.

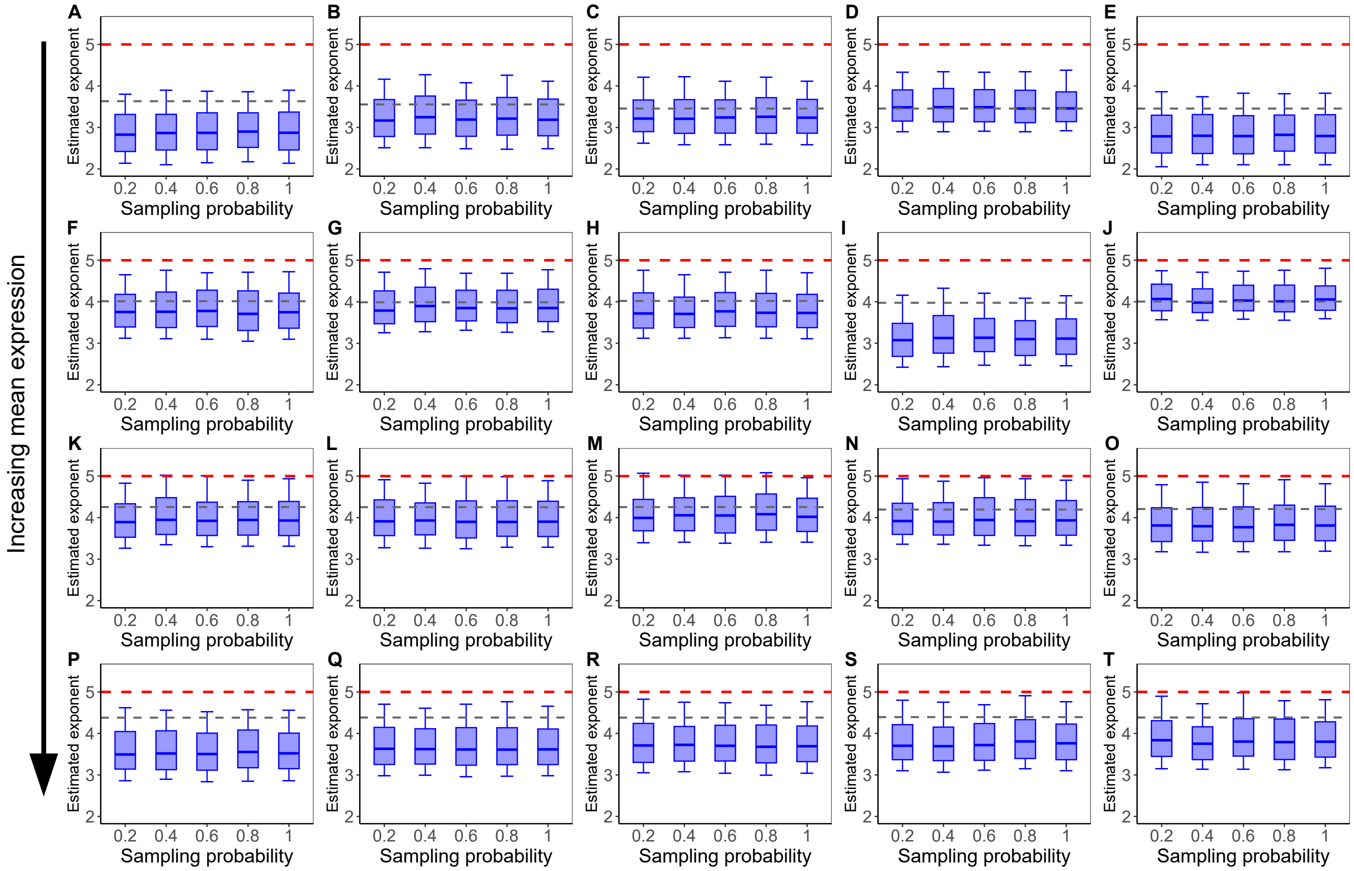

Fig. S8. Changing the mRNA capture success probability does little to affect the estimated exponent from linear regression on the mean mRNA count following induction. Panels A-T show the estimated exponent using linear regression from 20 different parameters sets using 5000 independent samples of the mean computed with sample size (number of cells) 1000, generated using the SSA. Each sample was resampled using binomial sampling procedure representing technical noise introduced by the mRNA capture procedure. The following binomial sampling probabilities of 0.2, 0.4, 0.6, 0.8, and 1.0 were used to represent various mRNA capture success rates. Boxplots show the median, IQR, and whiskers extended to the 10th and 90th percentiles. The grey dashed horizontal line shows the infinite sample size exponent, and the red line shows the theoretical short time exponent $(N - j + 1)$. In this case $N = 5, j = 1$. The distribution of estimated exponents remains the same as the sampling probability increases. Each row represents parameter sets whose mean at the final sample was approximately 1 (A-E), 5 (F-J), 8 (K-O), and 12 (P-T). Parameter sets were chosen from those randomly generated for Fig. 4. See SI Note 6.3 for details on the sampling procedure.

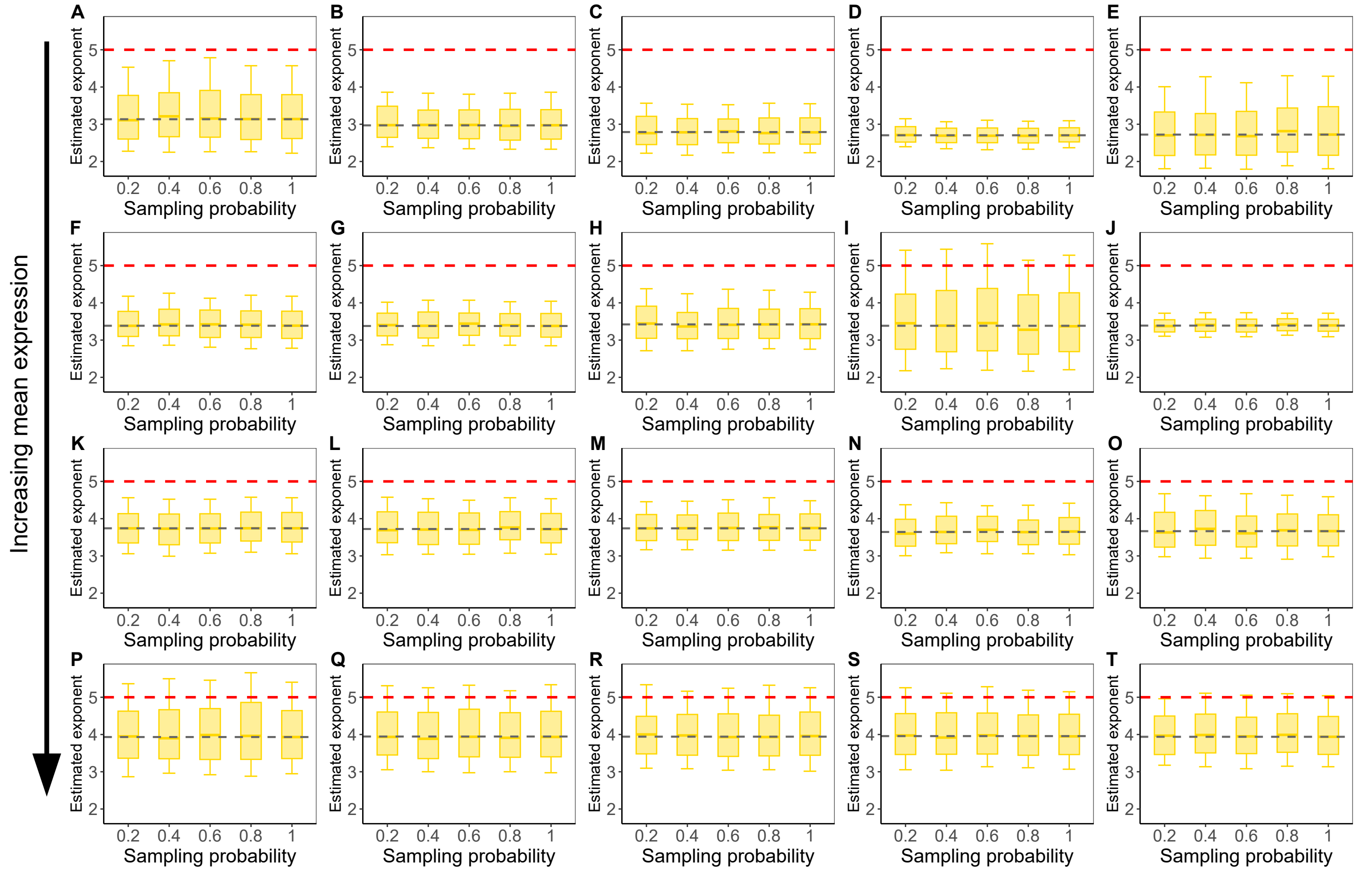

Fig. S9. Changing the mRNA capture success probability does little to affect the estimated exponent from non-linear regression on the mean mRNA count following induction. Panels A-T show the estimated exponent using linear regression from 20 different parameters sets using 5000 independent samples of the mean computed with sample size (number of cells) 1000, generated using the SSA. Each sample was resampled using binomial sampling procedure representing technical noise introduced by the mRNA capture procedure. The following binomial sampling probabilities of 0.2, 0.4, 0.6, 0.8, and 1.0 were used to represent various mRNA capture success rates. Boxplots show the median, IQR, and whiskers extended to the 10th and 90th percentiles. The grey dashed horizontal line shows the infinite sample size exponent, and the red line shows the theoretical short time exponent $(N - j + 1)$. In this case $N = 5, j = 1$. The distribution of estimated exponents remains the same as the sampling probability increases. Each row represents parameter sets whose mean at the final sample was approximately 1 (A-E), 5 (F-J), 8 (K-O), and 12 (P-T). Parameter sets were chosen from those randomly generated for Fig. 4. See SI Note 6.3 for details on the sampling procedure.

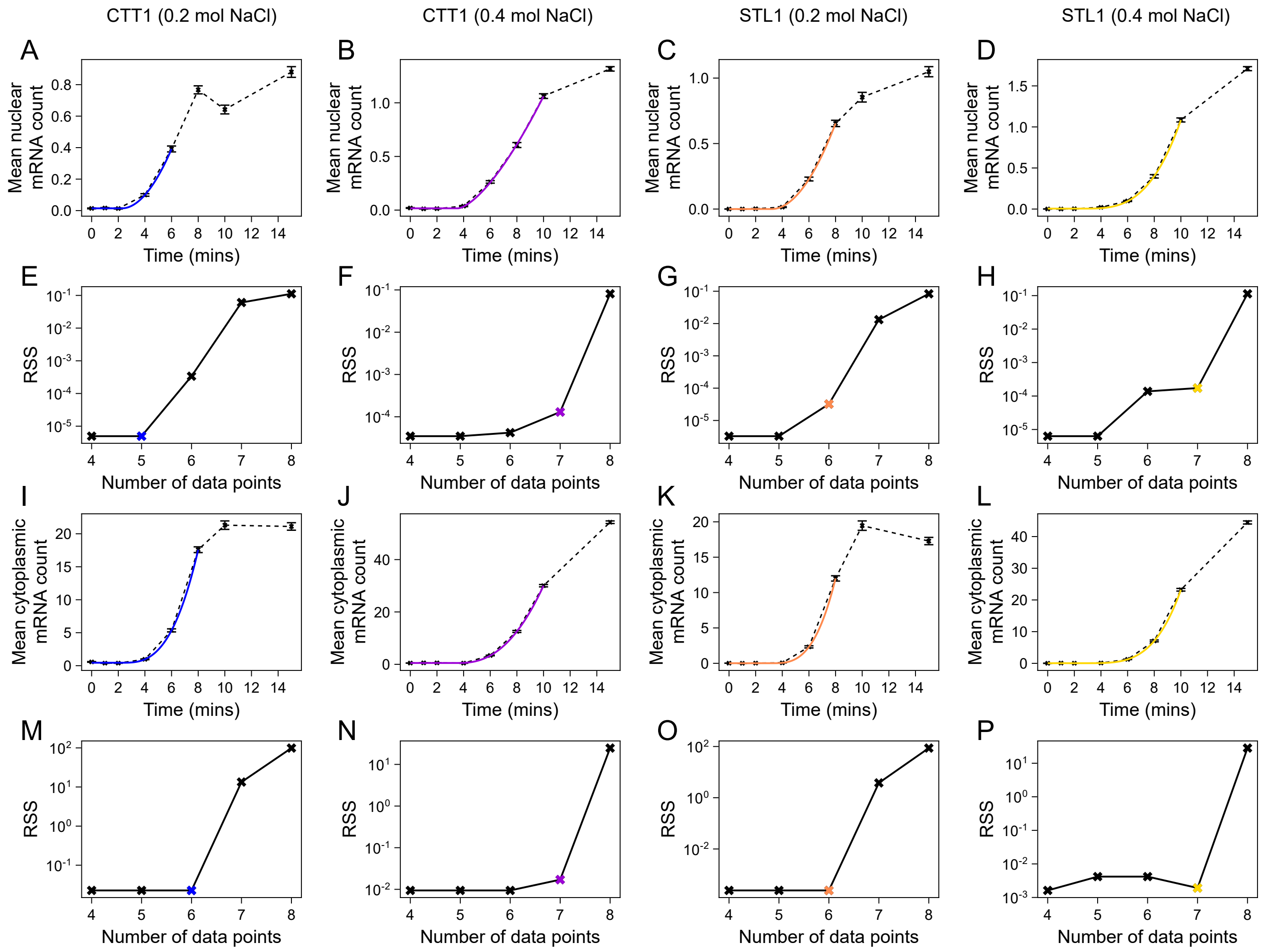

Fig. S10. Residual sum of squares (RSS) for each yeast data induction curve can be used to determine the optimal number of data points for regression. We choose the optimal number of data points as the maximum number of points that minimises the RSS. This can be determined as the value when the RSS significantly drops off from the right. The optimal number for each curve is marked by a coloured cross. Each RSS plot (below) is coupled with its corresponding data induction curve (above). Left to right: CTT1 0.2 mol NaCl, CTT1 0.4 mol NaCl, STL1 0.2 mol NaCl, STL1 0.4 mol NaCl. See SI Note 6.4 for further details.

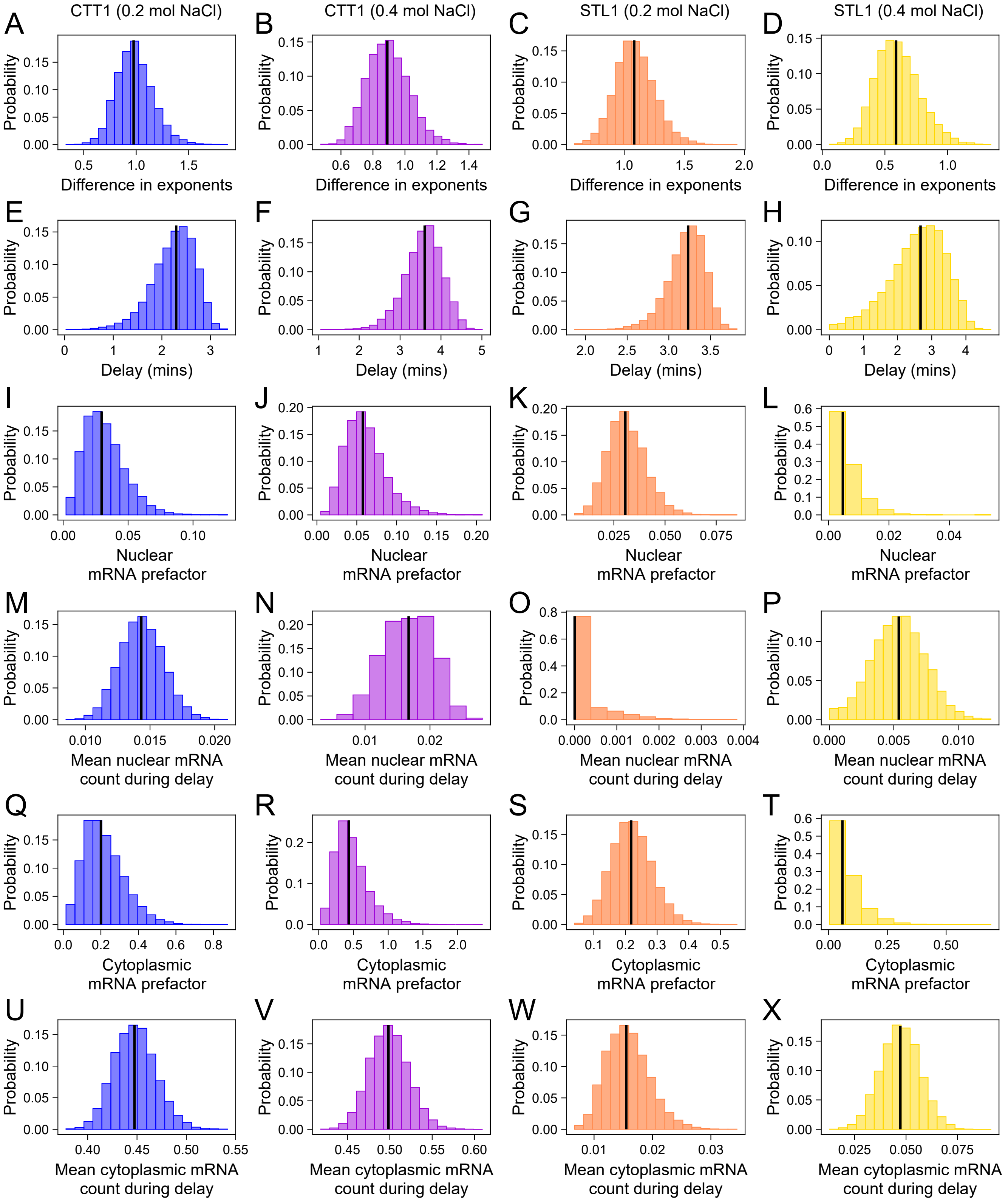

Fig. S11. Sampling distributions of estimated parameters from non-linear regression fitting of yeast mRNA induction curves. Left to right: CTT1 0.2 mol NaCl, CTT1 0.4 mol NaCl, STL1 0.2 mol NaCl, STL1 0.4 mol NaCl. (A-D) Distributions of the difference between each nuclear and cytoplasmic exponent from bootstrapping. (E-H) Distributions of the fixed time delay ($t_0$) before transcription initiation from bootstrapping. (I-L) Distributions of the nuclear mRNA prefactor from bootstrapping. (M-P) Distributions of the mean nuclear mRNA count during the delay from bootstrapping. (Q-T) Distributions of the cytoplasmic mRNA prefactor from bootstrapping. (U-X) Distributions of the mean cytoplasmic mRNA count during delay from bootstrapping. Sampling distributions were computed over 10,000 independent bootstrapped samples. The median of each distribution is marked by a vertical black line. See SI Note 6.4 for further details.

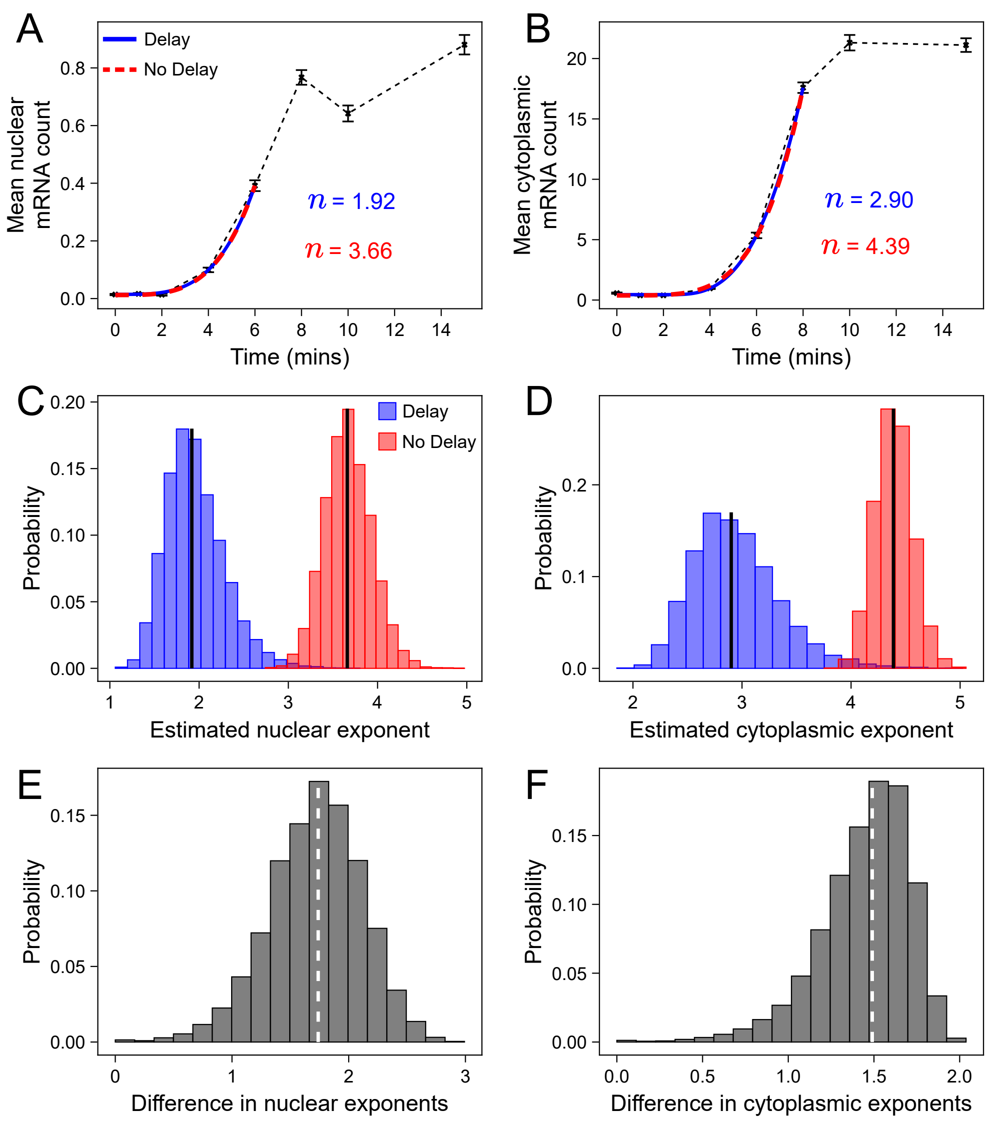

Fig. S12. Estimation of the exponent is sensitive to the time delay between the stimulus and the onset of transcription initiation. The exponent can be overestimated if the beginning of initiation is not estimated. (A-B) Fit of power law model with delay (blue, as seen in Fig. 5 C and K in the main text) and without delay (red) (given by Eqs. (87) and (53), respectively) to nuclear and cytoplasmic mean mRNA count curves (CTT1, 0.2 mol NaCl). (C-D) Distributions of estimated nuclear and cytoplasmic exponents from bootstrapped mean curves (vertical black line is the median). (E-F) Distributions of the difference between the nuclear (E) and cytoplasmic (F) exponents estimated from models with delay and no delay (vertical white dashed line is the median). Sampling distributions were computed over 10,000 independent bootstrapped samples. Both curves fit well to data but give different exponents whose sampling distributions do not significantly overlap. See SI Note 6.5 for further details.

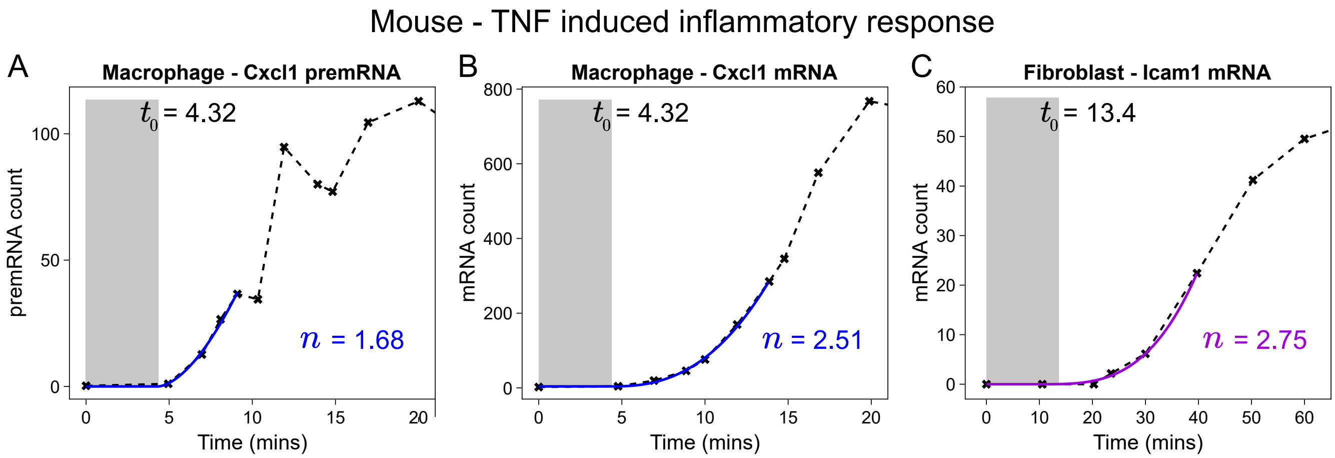

Fig. S13. RNA count induction curves of genes Cxcl1 (blue) and Icam1 (purple) in mouse macrophage and fibroblast cells under tumor necrosis factor (TNF) induced inflammatory response [3]. (A) Cxcl1 premRNA (unspliced mRNA) in macrophage cells. (B) Cxcl1 (spliced) mRNA in macrophage cells. (C) Icam1 (spliced) mRNA in fibroblast cells. Delay shown shaded in grey. The estimated exponent $n$ is shown next to the best fit curve. For Cxcl1 the mRNA curve was fit to the model given in Eq. (87) to infer the delay, which was fixed for the premRNA fit. Difference in exponents of 0.82 is consistent with splicing being described by an effective first-order reaction. See SI Note 6.6 for further details.

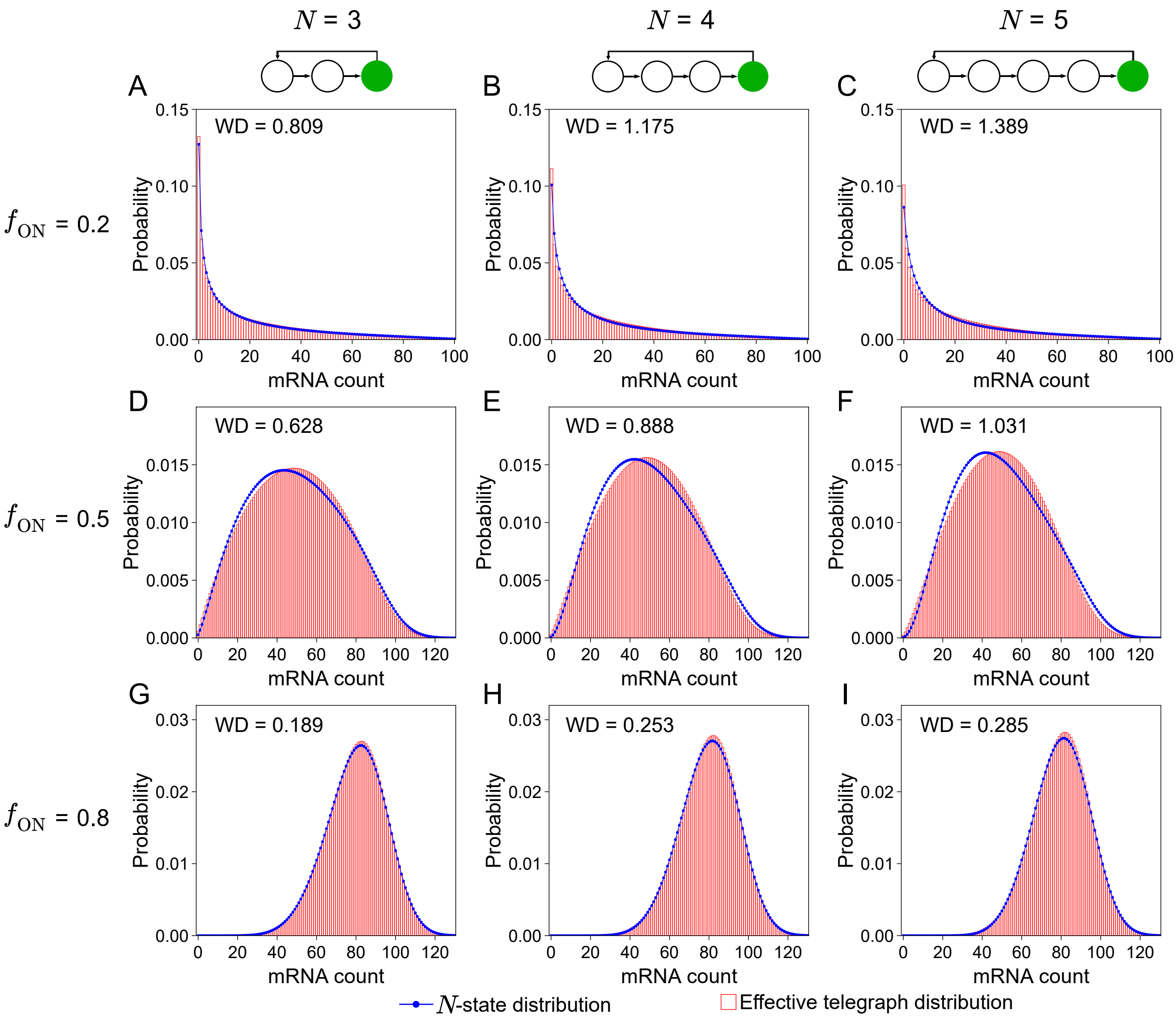

Fig. S14. Even models with identical gene state switching rates are, in steady-state, well approximated by the telegraph model. Steady-state distributions of the mRNA count for $N = 3$ (A, D, G), 4 (B, E, H), and 5 (C, F, I) state models and their effective telegraph distributions are indistinguishable despite the gene state switching rates, $k_i$ for $i = 1, \ldots, N-1$, being identical. As the number of rate-limiting steps increases, the Wasserstein distance (WD) between the $N$-state and effective telegraph distributions increases for each fixed mean fraction of time spent in the active state, $f_{ON}$, but the distributions remain qualitatively identical. The parameters $k_i$ for $i = 1, \ldots, N-1$ were chosen such that $k_i = (N-1)a$ where $a = 0.375$ (A, B, C), 1.5 (D, E, F), 6.0 (G, H, I). The other parameters were fixed for all distributions: $k_N = 1.5$, $\rho = 100$, and $d = 1$. This resulted in $f_{ON} = 0.2$, 0.5, and 0.8. All distributions are of shape II (unimodal with Fano factor $\geq 2$), as described in Fig. 2 in the main text. For more details see SI Note 6.7 and Table S8 for rate parameters.

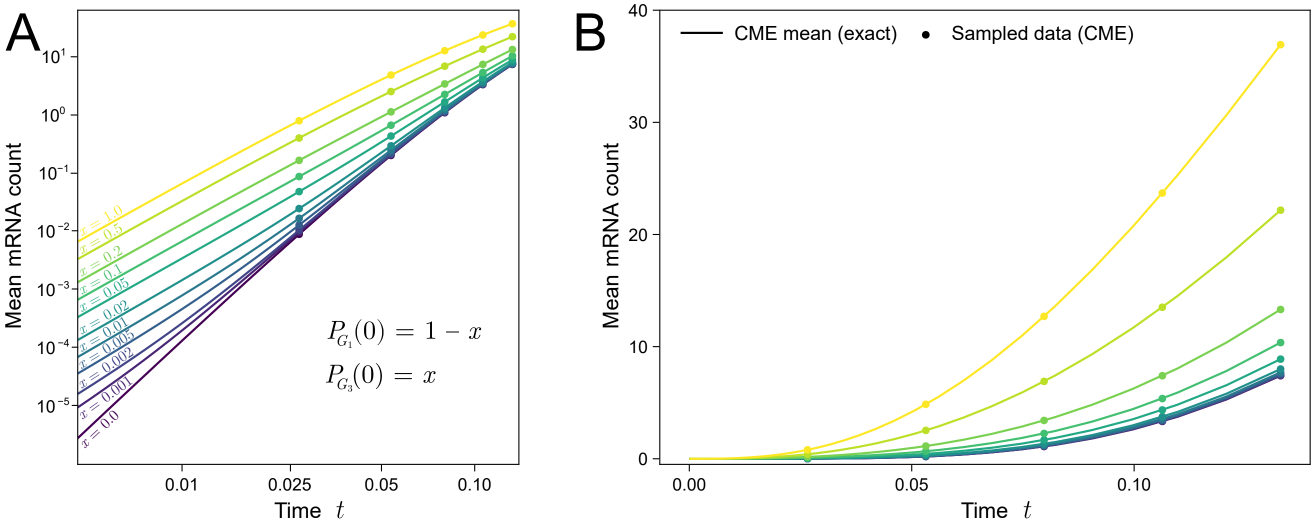

C

| Initial condition | | Regression | |
|---|---|---|---|
| $P_{G_1}(0)$ | $P_{G_3}(0)$ | $R^2$ | Estimated exponent |
| 0.0 | 1.0 | 0.999 | 2.28 |
| 0.5 | 0.5 | 1.00 | 2.36 |
| 0.8 | 0.2 | 0.997 | 2.52 |
| 0.9 | 0.1 | 0.998 | 3.18 |
| 1.0 | 0.0 | 1.00 | 4.63 |

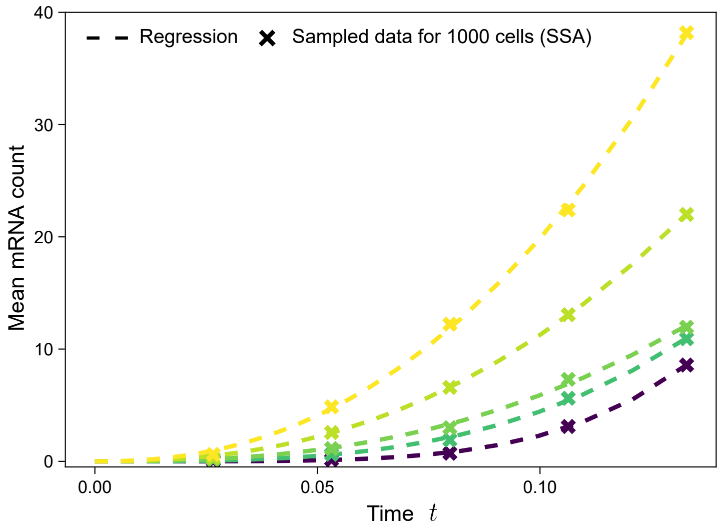

Fig. S15. The mean mRNA count versus time curve for short times is well approximated by a single power law with an exponent that is smaller than the total number of gene states even when the initial gene state distribution is mixed. Consider a 5-state model and assume that initially a gene is in state $G_1$ with probability $P_{G_1}(0) = 1 - x$ and in state $G_3$ with probability $P_{G_3}(0) = x$ where $x$ is varied between 0.0 and 1.0. (A-B) The mean mRNA count versus time for various different initial gene state probabilities ($x$) computed by direct numerical integration of the first moment equations of the CME for the 5-state model with the following rate parameters: $k_1 = 18.6, k_2 = 18.3, k_3 = 13.0, k_4 = 29.7, k_5 = 1.3, \rho = 880, d = 1.0$. (C) Results from non-linear regression fitting of the single power law model (see Eq. (53)) to five measurements of the mean mRNA count computed using the SSA with sample size (number of cells) 1000. Table (left) shows each initial condition, the $R^2$ of the regression model (3 s.f.), and the estimated exponent. The mean mRNA count plot (right) shows the sampled mean measurements (crosses) and their corresponding regressions (dashed lines). For more details see SI Note 6.8.

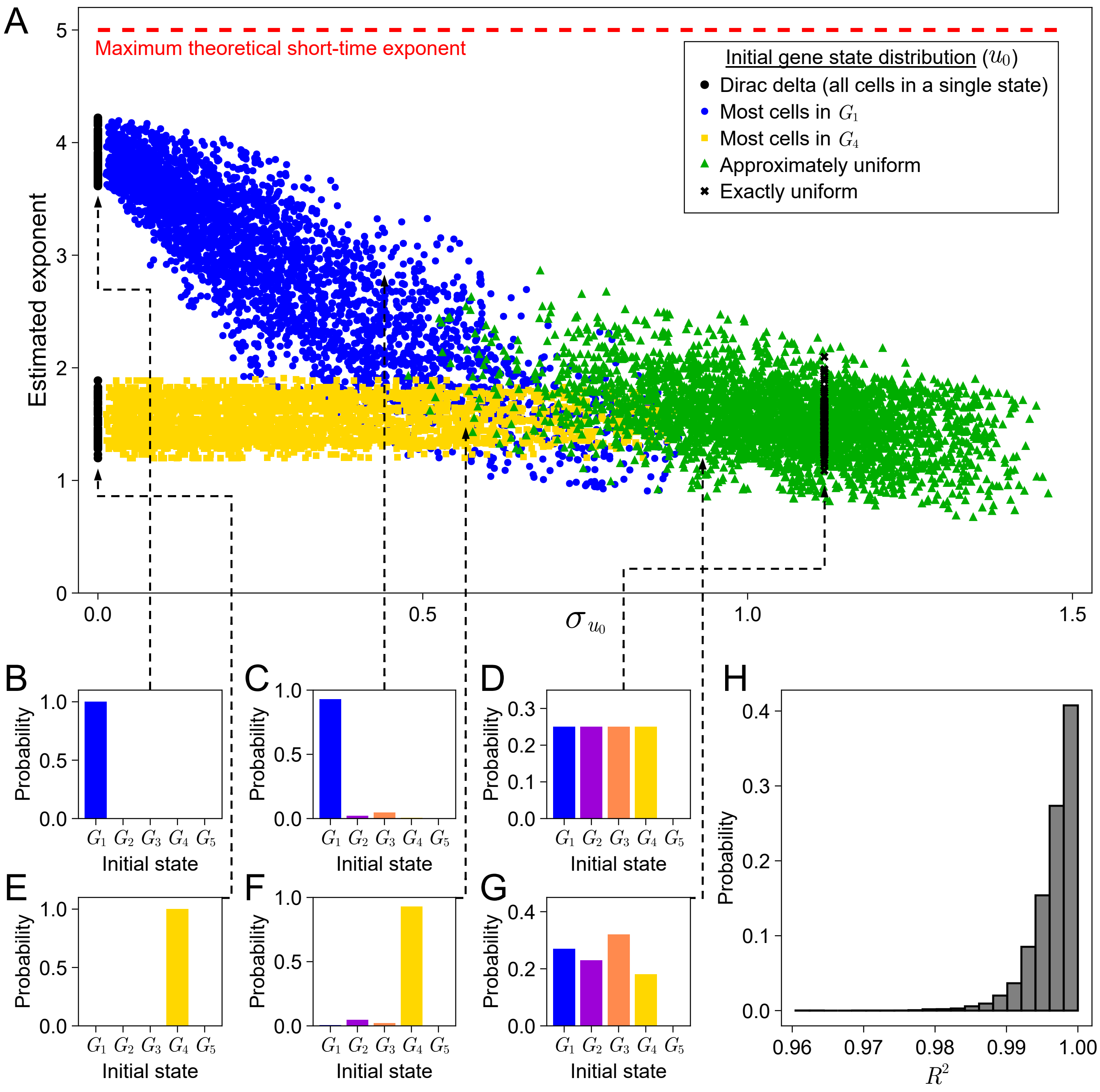

Fig. S16. Dependence of the short-time exponent of the 5-state model (following induction) on the standard deviation of the initial gene state distribution (A) and its type (B-G). The exponent is estimated by fitting a single power law to data generated by direct numerical integration of the moment equations of the CME. See SI Note 5.3 for further details. The following initial gene state distributions were used to sample various initial conditions: a Dirac delta distribution on states $G_1$ (B) and $G_4$ (E) (black circles); a distribution in which $P_{G_1}(t = 0) > 0.9$ i.e. most cells start in state $G_1$ (C) (blue circles); a distribution in which $P_{G_4}(t = 0) > 0.9$ i.e. most cells start in state $G_4$ (F) (yellow squares); an exact uniform distribution $P_{G_i}(t = 0) = 0.25$ for $i = 1, 2, 3, 4$ (D) (black crosses); and an approximate uniform distribution i.e. all inactive states are approximately equally probable (G) (green triangles). (H) The distribution of $R^2$ for all regressions performed for (A). Over 95% of regressions gave $R^2 \geq 0.99$, indicating very high goodness of fit by the single power law model across the parameter space and for all types of initial conditions. See SI Note 6.8 for further details on the parameter set and initial condition sampling procedures.
